# Missing-In-Metastasis / Metastasis Suppressor 1 regulates B cell receptor signaling, B cell metabolic potential and T cell-independent immune responses

**DOI:** 10.1101/782276

**Authors:** Alexey V Sarapulov, Petar Petrov, Sara Hernández-Pérez, Vid Šuštar, Elina Kuokkanen, Rufus Vinod, Marika Vainio, Lena Cords, Marco Fritzsche, Yolanda R Carrasco, Pieta K Mattila

## Abstract

Efficient generation of antibodies by B cells is one of the prerequisites of protective immunity. B cell activation by cognate antigens via B cell receptors (BCR), or pathogen-associated molecules through pattern-recognition receptors, such as Toll-like receptors (TLRs), leads to transcriptional and metabolic changes that ultimately transform B cells into antibody producing plasma cells or memory cells. BCR signaling and a number of steps downstream of it rely on coordinated action of cellular membranes and the actin cytoskeleton, tightly controlled by concerted action of multiple regulatory proteins, some of them exclusive to B cells. Here, we dissect the role of Missing-In-Metastasis (MIM), or Metastasis suppressor 1 (MTSS1), a cancer-associated membrane and actin cytoskeleton regulating protein, in B cell-mediated immunity by taking advantage of MIM knockout mouse strain. We show undisturbed B cell development and largely normal composition of B cell compartments in the periphery. Interestingly, we found that MIM^−/−^ B cells are defected in BCR signaling in response to surface-bound antigens but, on the other hand, show increased metabolic activity after stimulation with LPS or CpG. *In vivo*, MIM knockout animals exhibit impaired IgM antibody responses to immunization with T cell-independent antigen. This study provides the first comprehensive characterization of MIM in B cells, demonstrates its regulatory role for B cell-mediated immunity, as well as proposes new functions for MIM in tuning receptor signaling and cellular metabolism, processes which may also contribute to the poorly understood functions of MIM in cancer.

## 1 Introduction

Adaptive immune responses, such as efficient clearing of pathogens while maintaining the homeostasis of the host, depend on fine-tuned balance of various signals. Increasing evidence points towards an important role of the actin cytoskeleton and plasma membrane organization at the cross-roads of various signaling pathways orchestrating lymphocyte action (Mattila et al., 2016). In B cells, the actin cytoskeleton enables changes in cell morphology, required, for instance, during the formation of the immunological synapse (Kuokkanen et al., 2015). Interestingly, actin cytoskeleton and plasma membrane also potently regulate B cell receptor (BCR) signaling (Treanor et al., 2010; Mattila et al., 2013, 2016). A plethora of cytoskeletal regulator proteins enable the multifaceted roles of the actin cytoskeleton in living cells. Lymphocytes generally present very characteristic protein expression patterns and considering the specialized functions of these immune cells, it is not surprising that this also holds true for the regulators of the actin cytoskeleton. One best-known example of such protein is Wiscott-Aldrich syndrome (WAS) protein (WASp), a critical regulator of lymphocyte function and an activator of Arp2/3 actin filament (F-actin) nucleator complex (Bosticardo et al., 2009).

A highly conserved, cancer-associated protein linked to the regulation of both the actin cytoskeleton and the plasma membrane, Missing in Metastasis (MIM) or Metastasis Suppressor 1 (MTSS1), is highly expressed in spleen and particularly in B cells (Mattila et al., 2003; BioGPS.org portal). MIM belongs to a family of proteins with a characteristic inverse Bin, Amphiphysin, Rvs (I-BAR) domain, or IRSp53 and MIM homology domain (IMD), which binds and deforms cellular membranes (Mattila et al., 2007; Safari and Suetsugu, 2012). It also directly interacts with and regulates actin via its C-terminal WASp homology 2 (WH2) domain (Mattila et al., 2003; Woodings et al., 2003) and indirectly via interactions with other actin regulatory proteins, such as cortactin and Rac1 GTPase (Mattila et al., 2003, 2007; Lin et al., 2005; Cao et al., 2012). MIM has also been shown to regulate bone marrow and lymphoid cell trafficking presumably through regulation of CXCR4 internalization as seen in cancer cell lines (Yu et al., 2012; Zhan et al., 2016; Li et al., 2017; Zhao et al., 2019). Importantly, MIM has been linked to various cancers, either as a putative tumor metastasis suppressor, or promoter (Machesky and Johnston, 2007). Genetic alterations in *MIM/MTSS1* gene were found in 6% of sequenced cancer samples and, depending on the cancer type, both diminished or increased gene expression profiles are seen (Petrov et al., 2019). Regarding hematopoietic malignancies, MIM is upregulated, for example, in hairy cell and mantle cell lymphomas as well as chronic lymphocytic leukaemia (CLL). In CLL, interestingly, the good prognosis samples exhibit highest levels of MIM while the poor prognosis samples show lower MIM levels in comparison to good prognosis samples (Petrov et al., 2019). In mice, it has been reported that upon aging MIM knockout animals develop lymphomas resembling diffuse large B cell lymphoma (DLBCL) (Yu et al., 2012). Moreover, a degenerative kidney disease, potentially linked to impaired cell-cell junction formation, as well as a defected dendritic spine formation and neuronal alterations have been reported in MIM knockout mice (Saarikangas et al., 2011, 2015). These findings illustrate the complexity of MIM function, the basis of which remains enigmatic due to the lack of understanding about the molecular mechanisms and connected pathways. Despite the reported high expression in B cells and the association with hematopoietic malignancies, nothing is known about the role of MIM in activation of adaptive immune responses.

In this study, we took advantage of a MIM knockout mouse model (MIM^−/−^, MIM-KO) (Saarikangas et al., 2011) to explore the physiological role of MIM in B cell compartment, specifically in early B cell activation and mounting of the antibody responses. While we found no defects in B cell development, MIM-deficiency caused a variety of changes in mature B cells. MIM^−/−^ B cells showed significantly reduced signaling upon stimulation with surface-bound antigens mimicking activation via immunological synapse. T cell-independent IgM responses were reduced in MIM^−/−^ mice, while on the other hand, T cell-dependent immune responses appeared normal. Unlike BCR stimulation, MIM^−/−^ B cells were robustly activated by TLR agonists that, interestingly, also led to increased metabolic activity in cells lacking MIM. Our study highlights the complex role of MIM in different cellular functions and can serve as a stepping stone for unveiling the role of MIM in hematopoietic cancers.

## 2 Materials and Methods

### 2.1 Antibodies and chemicals

List of antibodies and reagents used in the study can be found in Table I.

**Table 1.** Key reagents table

| Name | Catalog # | Company | Application |
| --- | --- | --- | --- |
| AffiniPure Donkey Anti-Mouse IgM, $\mu$ Chain Specific | 715-005-020 | Jackson ImmunoResearch | Stimulatory antibodies |
| AffiniPure F(ab)<sub>2</sub> Fragment Donkey Anti-Mouse IgM | 715-006-020 | Jackson ImmunoResearch | Stimulatory antibodies |
| AF647 AffiniPure F(ab)<sub>2</sub> Fragment Donkey Anti-mouse IgM | 715-606-020 | Jackson ImmunoResearch | Stimulatory antibodies |
| Monobiotinylated Purified Rat anti-mouse Ig $\kappa$ light chain | 559749 or 21343 | BD Biosciences or Thermo Fisher Scientific | Stimulatory antibodies |
| Purified Rat Anti-Mouse IgM - Clone II/41 (RUO) | 553435 | BD Biosciences | ELISA, capture antibody |
| AffiniPure Goat Anti-Mouse IgG, Fcy Fragment Specific | 115-005-071 | Jackson ImmunoResearch | ELISA, capture antibody |
| Goat Anti-Mouse IgG1-BIOT | 1071-08 | SouthernBiotech | ELISA, detection antibody |
| Goat Anti-Mouse IgG2b-BIOT | 1091-08 | SouthernBiotech | ELISA, detection antibody |
| Goat Anti-Mouse IgG2c-BIOT | 1078-08 | SouthernBiotech | ELISA, detection antibody |
| Goat Anti-Mouse IgG3, Human/Bovine/Horse SP ads-BIOT | 1103-08 | SouthernBiotech | ELISA, detection antibody |
| Goat Anti-Mouse IgG Fc-BIOT | 1033-08 | SouthernBiotech | ELISA, detection antibody |
| Biotin Rat Anti-Mouse IgM, Clone R6-60.2 (RUO) | 553406 | BD Biosciences | ELISA, detection antibody |
| C57BL/6 Mouse Immunoglobulin Panel | 5300-01B | SouthernBiotech | ELISA, standard |
| Purified Rat Anti-Mouse CD16/CD32 | 553142 | BD Biosciences | Flow cytometry, Fc-block |
| FITC Rat Anti-Mouse IgG1 | 553443 | BD Biosciences | Flow cytometry, CSR |
| FITC Rat Anti-Mouse IgG2b | 553395 | BD Biosciences | Flow cytometry, CSR |
| FITC Rat Anti-Mouse IgG3 | 553403 | BD Biosciences | Flow cytometry, CSR |
| AffiniPure Goat anti-Mouse IgG2c, FITC-conjugated | 115-095-208 | Jackson ImmunoResearch | Flow cytometry, CSR |
| LPS, Lipopolysaccharides from Escherichia coli | L2887-5MG | Sigma-Aldrich | CSR, Metabolism |
| CpG ODN 1826 | tlrl-1826 | Invivogen | Metabolism |
| CD40L, Recombinant Mouse CD40 Ligand/TNFSF5 (HA-tag) | 8230-CL-050 | R&D Systems | CSR |
| IL-4, Recombinant Mouse IL-4 | 404-ML-010 | R&D Systems | CSR, Metabolism |
| IFNγ, Recombinant Mouse IFN-g | 575304 | BioLegend | CSR |
| TGF-β, Recombinant Mouse TGF-beta 1 | 7666-MB-005 | R&D Systems | CSR |
| BD Horizon™ V450 Rat Anti-Mouse IgM Clone R6-60.2 | 560575 | BD Biosciences | Flow cytometry |
| AF488 anti-mouse IgD [11-26c.2a] | 405718 | Biolegend | Flow cytometry |
| AF700 anti-mouse CD19 [6D5] | 115528 | Biolegend | Flow cytometry |
| APC anti-mouse CD19 [6D5] | 115512 | Biolegend | Flow cytometry |
| APC/Cy7 anti-mouse/human CD45R/B220 [RA3-6B2] | 103224 | Biolegend | Flow cytometry |
| APC anti-mouse CD21/CD35 (CR2/CR1) [7E9] | 123412 | Biolegend | Flow cytometry |
| Alexa Fluor® 594 anti-mouse CD23 [B3B4] | 101628 | Biolegend | Flow cytometry |
| PE anti-mouse CD93 (AA4.1, early B lineage) [AA4.1] | 136503 | Biolegend | Flow cytometry |
| PE anti-mouse CD3e [145-2C11] | 100308 | Biolegend | Flow cytometry |
| FITC anti-mouse CD4 [GK1.5] | 100406 | Biolegend | Flow cytometry |
| APC anti-mouse CD5 [53-7.3] | 100626 | Biolegend | Flow cytometry |
| Tom20 (FL-145) | sc-11415 | Santa Cruz Biotechnology | Flow cytometry, metabolism |
| Goat anti-Rabbit IgG (H+L), AF488 | A-11008 | Thermo Fisher Scientific | Flow cytometry, metabolism |
| MTSS1 (P549) Ab | 4385S | Cell Signaling Technologies | Immunoblotting |
| MTSS1 (N747) Ab | 4386S | Cell Signaling Technologies | Immunoblotting |
| Phospho-Zap-70 (Tyr319)/Syk (Tyr352) Ab | 2701P | Cell Signaling Technologies | Immunoblotting, |

|  |  |  | microscopy |
| --- | --- | --- | --- |
| Syk (D3Z1E) XPγ Rabbit mAb | 13198S | Cell Signaling Technologies | Immunoblotting |
| Phospho-Lyn (Tyr507) Ab | 2731P | Cell Signaling Technologies | Immunoblotting |
| Lyn (C13F9) rabbit mAb | 2796S | Cell Signaling Technologies | Immunoblotting |
| Phospho-CD19 (Tyr531) Ab | 3571S | Cell Signaling Technologies | Immunoblotting |
| CD19 Ab | 3574S | Cell Signaling Technologies | Immunoblotting |
| Phospho-PI3 Kinase p85 (Tyr458)/p55 (Tyr199) Ab | 4228S | Cell Signaling Technologies | Immunoblotting |
| Phospho-CD19 (Tyr531) Ab | 3571S | Cell Signaling Technologies | Immunoblotting |
| Phospho-Akt (Ser473) (193H12) Rabbit mAb | 4058S | Cell Signaling Technologies | Immunoblotting |
| Akt1 (C73H10) Rabbit mAb | 2938S | Cell Signaling Technologies | Immunoblotting |
| Phospho-NF-kappa-B p65 (Ser536) (93H1) Rabbit mAb | 3033S | Cell Signaling Technologies | Immunoblotting |
| NF-κB p65 (D14E12) XP$\otimes$ Rabbit mAb | 8242S | Cell Signaling Technologies | Immunoblotting |
| p44/42 MAPK (Erk1/2) Ab | 9102S | Cell Signaling Technologies | Immunoblotting |
| Phospho-p44/42 MAPK (Erk1/2) (Thr202/Tyr204) Ab | 9101S | Cell Signaling Technologies | Immunoblotting |
| Phospho-Btk (Tyr223) (D1D2Z) Rabbit mAb | 87457 | Cell Signaling Technologies | Immunoblotting |
| Peroxidase AffiniPure Goat Anti-Rabbit IgG (H+L) | 111-035-144 | Jackson ImmunoResearch | Immunoblotting |
| Peroxidase AffiniPure Goat Anti-Mouse IgG, Fcy Specific Ab | 115-035-071 | Jackson ImmunoResearch | Immunoblotting |
| Donkey Anti-Rabbit IgG Ab, IRDye 800CW Conjugated | 926-32213 | LI-COR Biosciences | Immunoblotting |
| AF555 Phalloidin | A34055 | Thermo Fisher Scientific | Microscopy |
| AF647 anti-mouse/human CD45R/B220 Ab | 103229 | BioLegend | Microscopy |
| Anti-Phosphotyrosine Ab, clone 4G10$\otimes$ | 05-321 | Merck Millipore | Microscopy |
| FITC Rat Anti-Mouse IgG2b | 553395 | BD | Microscopy |
| CXCL13 | 250-24-5ug | PeproTech | Microscopy, SLB |
| AlexaFluor 488 Mouse Anti-Btk (pY223)/Itk (pY180) | 564847 | BD Biosciences | Microscopy |
| Donkey anti-rabbit IgG, AF488 conjugate | A-21206 | Thermo Fisher Scientific | Microscopy |

### 2.2 Mice

MIM knockout mouse colony was a kind gift from Prof. Pekka Lappalainen and Dr. Pirta Hotulainen from the University of Helsinki and Minerva Foundation Institute for Medical Research (Saarikangas et al., 2011). The strain, in C57Bl/6 background, had no apparent health problems until the age of 8 months when mice were latest sacrificed, however, we observed that from all genotyped animals that were kept alive, 18 pups developed hydrocephaly over the study period. Among them, 17 were knockout, 1 heterozygote, and none wild-type. To generate this strain, Saarikangas et al (Saarikangas et al., 2011) introduced a *Neo*-cassette, containing several stop codons, by homologous recombination into Exon 1 of *MIM*/*Mtss1* gene in 129/Sv ES-cells. Chimeric mice were backcrossed to C57Bl/6J background for several generations and the colony in Turku was established by breedings of heterozygote founder animals. All experiments were done with age- and sex-matched animals and WT littermate controls were used whenever possible.

### 2.3 Immunizations

At the age of 3-4 months, groups of WT and *MIM*^−/−^ females were immunized with NP_40_‑FICOLL (F-1420, Biosearch Technologies) for T-independent (TI) immunization or NP_31_‑KLH (N-5060, Biosearch Technologies) for T‑dependent (TD) immunization. Each mouse received 50 μg of antigen in 150 μL of PBS (NP_40_‑FICOLL) or PBS/Alum (77161, Thermo Fisher) adjuvant (2:1 ratio) (NP_31_‑KLH) solution by intraperitoneal injection. Blood (~100 μL) was sampled from lateral saphenous veins on day −1 (preimmunization) and every week after immunization on days +7, +14, +21 and +28 for both FICOLL and KLH cohorts. Secondary immunization of KLH cohort was performed on day +135 (0) and blood was sampled on days +134 (−1), +139 (+4), +143 (+8) and +150 (+15). Coagulated blood was spun at +4°C / 2000 rpm for 10 min and serum was collected and stored at −20°C.

All animal experiments were approved by the Ethical Committee for Animal Experimentation in Finland. They were done in adherence with the rules and regulations of the Finnish Act on Animal Experimentation (62/2006) and were performed according to the 3R-principle (animal license numbers: 7574/04.10.07/2014, KEK/2014-1407-Mattila, 10727/2018).

### 2.4 ELISA

Total and NP-specific antibody levels were measured by ELISA on half-area 96-well plates (Greiner Bio-One, 675061). Wells were coated overnight at +4°C with capture antibodies (2 μg/mL) or NP-conjugated carrier proteins, NP_(1-9)_-BSA or NP_(>20)_-BSA (N-5050L, N-5050H, Biosearch Technologies) at 50 μg/mL in 25 μL PBS. Non-specific binding sites were blocked for 2 hours in 150 μL of blocking buffer (PBS, 1% BSA, 0.05% NaN_3_). Appropriate, experimentally determined dilutions (see below) of 50 μL serum samples in blocking buffer were added for overnight incubation at +4°C. Biotin-conjugated detection antibodies (2 μg/mL) in 50 μL of blocking buffer were added for 1 hour followed by 50 μL ExtrAvidin-Alkaline phosphatase (E2636, Sigma-Aldrich, 1:5000 dilution) in blocking buffer for 1 hour at room temperature (RT). In between all incubation steps, plates were washed with 150 μL washing buffer (PBS, 0.05% Tween-20) either 3 times for the steps before sample addition or 6 times after addition of the mouse sera. The final wash was completed with 2 times wash with 150 μL of water. Finally, 50 μL of Alkaline phosphatase-substrate, SIGMAFAST p-Nitrophenyl phosphate, (N2770, Sigma-Aldrich) solution was added and OD was measured at 405 nm. Serum dilutions were determined experimentally to fall into linear part of the dose-response curve of the absorbance measurements for any given isotype and typical values are as follows: IgM levels (1:3000– 1:4000), IgG levels (1:20000–1:80000). Different dilutions of AP-streptavidin were used where necessary. Typical time for AP-substrate incubation before measurement was about 30 min at RT.

All ELISA samples were run in duplicates, OD values were averaged and blank background was subtracted. Absolute concentrations of total antibody levels were extrapolated from calibration curves prepared by serial dilution of mouse IgM or subclasses of IgG from C57Bl/6 immunoglobulin panel. Relative NP-specific antibody levels were extrapolated from reference curves prepared by serial dilution of pooled serum, in which the highest dilution step received an arbitrary unit of 0.5.

### 2.5 Immunophenotyping

All cells were isolated in B cell isolation buffer (PBS, 2% FCS, 1 mM EDTA). Bone marrow cells were isolated by flushing the buffer through mouse femoral and tibial bones. Splenocytes were isolated by mashing the spleen in small buffer volumes with syringe plunger in 48-well plates. Peritoneal cavity cells were isolated by filling the cavity with ~10 mL buffer volume through puncture and collecting the fluid back. Cell suspensions were filtered through 70 μm nylon cell strainers. As a general flow cytometry protocol all following steps were done in flow cytometry buffer I (PBS, 1% BSA). Fc-block was done with 0.5 μL anti-mouse CD16/32 antibodies in 70 μL of flow cytometry buffer I for 10 min and cells were stained for 30 min. Washings were done 3 times in 150 μL of flow cytometry buffer I. All steps were carried out on ice in U-bottom 96-well plates at cell density of 0.25–0.5 × 10^6^/well. Before acquisition, cells were resuspended in 130 μL flow cytometry buffer II (PBS, 2.5% FCS). Samples were acquired on BD LSR Fortessa, equipped with four laser lines (405 nm, 488 nm, 561 nm, 640 nm). Compensation matrix was calculated and applied to samples either in BD FACSDiva™ software (BD Biosciences) or in FlowJo (Tree Star, Inc) based on fluorescence of conjugated antibodies using compensation beads (01-1111-41, Thermo Fisher Scientific). FMO (fluorescence minus one) controls were used to assist gating. Data was analyzed with FlowJo software.

### 2.6 B cell isolation

Splenic B cells were Isolated with EasySep™ Mouse B Cell Isolation Kit (19854, Stem Cells Technologies) according to manufacturer's instructions and let to recover in RPMI (10% FCS, 20 mM HEPES, 50 μM β-mercaptoethanol, 1:200 Pen/Strep) in an incubator at +37°C and 5% CO_2_ for 1-2 hours.

### 2.7 Class-switch recombination and proliferation

Isolated splenic B cells (~10–20×10^6^ cells) were stained first with 5 μL (5mM) Cell Trace Violet (C34557, Thermo Fisher Scientific) in 10 mL of PBS for 10 min at RT and let to recover in complete RPMI (+37°C, 5% CO_2_) for 1–2 hours. To induce class-switching, B cells were cultured on 24-well plates at 0.5×10^6^/mL density in complete RPMI supplemented with indicated doses of LPS (4 μg/mL), CD40L (150 ng/mL), IL-4 (5 ng/mL), IFN-γ (100 ng/mL) and TGF-β (3 ng/mL) for 3 days. Cells were blocked with anti-mouse anti‑CD16/32 and stained for 30 min with antibodies against IgG subclasses. Additionally, cells were stained with 4 μg/mL 7-AAD (ABD-17501, Biomol) for live/dead cell discrimination and samples were acquired on BD LSR II equipped with 3 laser lines (405 nm, 488 nm, 640 nm) and analyzed with FlowJo software.

### 2.8 B cell receptor signaling and immunoblotting

For analysis of BCR signaling, isolated splenic B cells were starved for 10 min in plain RPMI and 0.5 × 10^6^ cells in 100 μL of plain RPMI were stimulated in duplicates with anti-mouse IgM μ-chain-specific (anti-IgM) antibodies either in solution or bound to the culture dish surface, for 3, 7 and 15 min. For solution stimulation, 5 μg/mL of anti-IgM was used, in 96-well plates. For surface-bound mode, 48-well plates were coated with 5 μg/mL of anti-IgM antibodies in 120 μL of PBS at +4°C, overnight, and washed 3 times with 500 μL of ice-cold PBS before experiment. After activation, B cells were instantly lysed with 25 μL of 5x SDS lysis buffer (final: 62.5 mM TrisHCl pH ~6.8, 2% SDS, 10% Glycerol, 100mM β-mercaptoethanol, bromphenol blue) and sonicated for 7.5 min (1.5 mL tubes, high power, 30 s on/off cycle, Bioruptor plus, Diagenode). Lysates (20–30 μL) were run on 8–10% polyacrylamide gels and transferred to PVDF membranes (Trans-Blot Turbo Transfer System, Bio-Rad). Membranes were blocked with 5% BSA in TBS (TBS, pH ~7.4) for 1 hour and incubated with primary antibodies (~1:1000) in 5% BSA in TBST (TBS, 0.05% Tween-20) at +4°C, overnight. Secondary antibody incubations (1:20000) were done for 2 hours at RT in 5% milk in TBST for HRP-conjugated antibodies and with addition of 0.01% SDS for fluorescently-conjugated antibodies. Washing steps were done in 10 mL of TBST for 5 × 5 min. Membranes were scanned with Odyssey CLx (LI-COR) or visualized with Immobilon Western Chemiluminescent HRP Substrate (WBKLS0500, Millipore) and ChemiDoc MP Imaging System (Bio-Rad). Phospho-antibodies were stripped in 25 mM Glycine-HCl buffer, pH ~2.5 for 10 min, membranes were blocked and probed again for evaluation of total protein levels. Images were background subtracted and the raw integrated densities for each band were measured in ImageJ. Ratios of phosphorylated-vs-total protein levels were analyzed with ratio paired t test. For data presentation these ratios were normalized to WT value at 0 min.

### 2.9 Intracellular Ca^2+^ flux

Splenic B cells were resuspended at a concentration of 2.5-5 × 10^6^ cell/mL in RPMI supplemented with 20 mM HEPES and 2.5% FCS and loaded with 1 μM Fluo-4 (F14201, Thermo Fisher Scientific) and 3 μM Fura Red (F3021, Thermo Fisher Scientific) for 45 min (+37°C, 5% CO_2_). Cell suspension was diluted in 10 volumes of complete RPMI and incubated for 10-15 min at RT. Cells were centrifuged at 200 g, at RT for 5 min and resuspended at 2.5 × 10^6^ cells/mL in PBS supplemented with 20 mM HEPES, 5 mM glucose, 0.025% BSA, 1 mM CaCl_2_, 0.25 mM sulfinpyrazone (S9509, Sigma-Aldrich), 2.5% FCS. Cells were allowed to rest at RT for 20 min and were kept on ice before acquisition. Anti-IgM antibodies were added into prewarmed (+37°C, 5 min) B cell suspension aliquots to final concentrations of 10, 5, 2.5 and 1 μg/mL and samples were acquired on BD LSR Fortessa. Alternatively, equimolar concentrations of anti-IgM F(ab’)_2_ fragment were used. Fluorescence of Fluo-4 and Fura Red were recorded by a continuous flow for 5 min. Data was analyzed in FlowJo and presented as ratiometric measurement of Fluo-4/Fura Red median intensity levels.

Peritoneal cavity B cells were washed in L-15 medium, resuspended in 75 μL acquisition buffer (HBS (HEPES buffered saline):L-15 (1:1 ratio), 2.5 μM probenecid (P8761, Sigma-Aldrich)) and labeled by addition of 75 μL acquisition buffer with 10 μM Fluo-4 for 5 min at +37°C. Cells were washed in 1 ml, resuspended in 200 μL and divided into two wells. B cells were prestained for 10 min on ice with anti-CD23-AF594 antibodies, washed and resuspended in 100 μL of acquisition buffer on ice. Samples were prewarmed (+37°C) in a total volume of 300 μL of acquisition buffer and 50 μL of anti-IgM F(ab’)_2_-AF633 were added. Cells were acquired on BD LSR Fortessa for 3-5 min and analyzed in FlowJo.

### 2.10 Scanning electron microscopy

For the analysis of resting B cells, wells of the microscopy slides (10028210, Thermo Fisher Scientific) were coated with CellTak (354240, Corning) in PBS (3.5 μg/cm^2^ of surface area, according to manufacturer’s recommendations) for 20 min (RT), washed once with water and allowed to dry. For the analysis of activated B cells, wells were coated with 5 μg/mL of anti-IgM in PBS for 1 h (RT) and washed in PBS. 10^5^ B cells in 20 μL of complete RPMI were placed on coated wells for 10 min (+37°C, 5% CO_2_) and fixed by adding 20 μL PFA in PBS (4% PFA final, pH 7.0–7.5) for 15 min. Samples were further fixed in 4% PFA/2.5% gluteraldehyde in PBS for 30 min, washed in PBS and post-fixed in 1% OsO_4_ containing 1.5% potassium ferrocyanide, and dehydrated with a series of increasing ethanol concentrations (30%, 50%, 70%, 80%, 90%, 96% and twice 100%). Specimens were immersed in hexamethyldisilazane and left to dry by solvent evaporation. The cells were coated with carbon using Emscope TB 500 Temcarb carbon evaporator and imaged with Leo 1530 Gemini scanning electron microscope.

### 2.11 Immunofluorescence microscopy and cell spreading

#### TIRF microscopy

MatTek microscopy dishes were coated with 7.5 μg/mL of anti-IgM antibodies in PBS at +37°C for 30 min and washed once with PBS. Isolated splenic B cells (10^6^) were left unstained or labeled with 0.17 μL of anti-B220-AF647 antibodies in 400 μL PBS for 10 min in 1.5 mL tubes on ice, spun (2500 rpm, 5 min), washed twice in 900 μL of ice-cold PBS and resuspended in 200 μL of Imaging buffer (PBS, 10% FBS, 5.5 mM D-glucose, 0.5 mM CaCl_2_, 0.2 mM MgCl_2_). Equal amounts of unstained and labeled cells of different genotypes were mixed and loaded onto coated MatTek dishes at 35 μL/well. Cells were incubated for 10 min (+37°C, 5% CO_2_), fixed in pre-warmed (+37°C) 4% formaldehyde/PBS for 10 min (RT), permeabilized in 0.1% Triton X-100/PBS for 5 min (RT), washed once with PBS and blocked in blocking buffer (PBS, 1% BSA) at +4°C (overnight). Cells were stained with 1:50 Phalloidin-AF555 and 1:500 anti-pTyr primary antibody (4G10) in blocking buffer for 1 hr (RT), washed 4 times with PBS, and stained with 1:500 secondary anti-mouse IgG2b-AF488 in blocking buffer for 1 hr (RT), washed 4 times in PBS and imaged in PBS with total internal reflection fluorescence (TIRF) mode in DeltaVision OMX Imaging System (GE Healthcare). TIRF images of cortical actin and pTyr were processed with ImageJ macro using B220 and bright-field channels to discriminate between attached WT or MIM-KO cells. Spreading area (determined on pTyr channel), mean fluorescence intensity and total fluorescence intensity (integrated density) of phalloidin and pTyr staining of each cell were analyzed (~50–340 cells per sample). For cumulative scatter plots, equal numbers (here 92 cells) were randomly selected from each experiment.

#### Spinning disk confocal microscopy

12-well PTFE diagnostic slides (Thermo Fisher Scientific, #10028210) were coated with 5 μg/mL anti-mouse IgM (μ-chain-specific antibodies) in PBS at +4°C O/N and washed with PBS. As a non-activated control, wells were coated with 4 μg/mL fibronectin. Isolated splenic B cells (10^6^/ml) were labeled with 1μM CFSE (21888, Sigma-Aldrich) or left unlabeled. Equal amounts of unstained and labeled cells of different genotypes were mixed (1:1 ratio) and seeded at density of 100.000 cells/well. Dye-switched experiments were performed systematically. Cells were incubated for 3, 5, 7, 10 or 15 min (+37°C, 5% CO_2_), fixed in 4% formaldehyde/PBS for 10 min (RT) and permeabilized/blocked in 0.3% Triton X-100/5% donkey serum/PBS for 20 min (RT). Staining was performed in 0.3% Triton X-100/1% BSA/PBS at +4°C O/N, followed by washes with PBS and incubation with the secondary antibodies for 30 min at room temperature in PBS. Samples were mounted in FluoroMount-G (Thermo Fisher Scientific). Images were acquired on 3i CSU-W1 (Intelligent Imaging Innovations) Marianas spinning disk confocal microscope equipped with 63x Zeiss Plan-Apochromat objective and a Photometrics Prime BSI sCMOS camera. Cells were visualized at the plane of contact and 5-10 fields of view per sample were acquired. Images of F-actin, pBtk and pSyk were processed with ImageJ using CTV channel to discriminate between WT or MIM-KO cells. Spreading area (determined on the phalloidin channel) and mean fluorescence intensity of pBtk or pSyk staining per cell were analyzed (~25–150 cells per condition per experiment).

### 2.12 Supported lipid bilayers

Artificial planar lipid bilayers containing GPI-linked mouse ICAM-1 (200 molecules/μm^2^) were formed as previously described (Grakoui et al., 1999; Carrasco et al., 2004). Briefly, unlabeled GPI-linked ICAM-1 liposomes and liposomes containing biotinylated lipids were mixed with 1,2-dioleoyl-PC (DOPC) (850375P, Avanti lipids, Inc) at various ratios to obtain specified molecular densities. Planar membranes were assembled on FCS2 dosed chambers (Bioptechs) and blocked with PBS/2% FCS for 1 h at RT. Antigen was tethered by incubating membranes with AF647-streptavidin, followed by monobiotinylated anti-kappa light chain antibodies (20 molecules/μm^2^). The isolated B cells from WT and MIM^−/−^ mice were labeled with 1μM CFSE (21888, Sigma-Aldrich) or left unlabeled, mixed at 1:1 ratio, and injected into prewarmed chambers (4 × 10^6^ cells/chamber, +37°C) with 100 nM recombinant murine CXCL13. Fluorescence, differential interference contrast (DIC), and interference reflection microscopy (IRM) images were acquired at the plane of the cell contact in one position once every 30 s immediately after injecting the cells into the chamber for 10 min. After the 10 min movie was acquired, several snapshots were acquired for quantification of the mature synapses in different locations of the chamber at 10-15 min after cell injection. All assays were performed in PBS, supplemented with 0.5% FCS, 0.5 g/L D-glucose, 2 mM MgCl and 0.5 mM CaCl_2_. Images were acquired on Zeiss Axiovert LSM 510-META inverted microscope, equipped with 40x oil-immersion objective (Madrid), or Zeiss LSM 780 inverted microscope, equipped with 40x water-immersion objective (Turku), and analyzed by ImageJ. The spreading area (determined on IRM channel), area of collected antigen and mean fluorescence intensity of antigen were quantified from each experiment (~100 cells per experiment).

### 2.13 Intracellular Ca^2+^ flux on supported lipid bilayers

Splenic WT or MIM^−/−^ B cells (3.2 × 10^6^) were resuspended in 75 μL of L-15 medium and labeled by addition of 75 μL of HBS (HEPES buffered saline), supplemented with 2.5 μM probenecid and 20 μM Fluo4 for 5 minutes at +37°C. Cells were washed in 1 mL HBS-probenecid and resuspended in 500 μL HBS-probenecid for immediate injection into FCS2 chambers. Acquired movies were preprocessed with ImageJ and analyzed with a MATLAB implemented high-throughput software *CalQuo^2^* (Lee et al., 2017). Cells were categorized as single peak, oscillatory or not triggering. Cells showing more than two intensity peaks are classified as oscillatory. Data presented as mean percentages of 3 independent experiments with at least 1000 cells analyzed per experiment.

### 2.14 Metabolic assay

Splenic B cells were seeded at a density of 10^6^ cells/mL in complete RPMI and treated with indicated combinations of IL-4 (10 ng/mL), anti-mouse IgM (10 μg/mL), LPS (4 μg/mL) and CpG (10 μg/mL) for 24 h at +37°C, 5% CO_2_ in a humidified incubator. Cells were then spun and resuspended in Seahorse XF RPMI (103576-100, Agilent), supplemented with 1 mM pyruvate, 2 mM L-glutamine and 10 mM D-glucose. Cell number was adjusted and 0.15×10^6^ cells were seeded per well on a 96-well XF plate, pre-coated with CellTak (354240, Corning). Plate coating was done with 22.4 μg/mL CellTak in NaHCO_3_, pH 8.0 at +4°C overnight, followed by two washings with water. Seeded cells were spun at 200 g for 1 min with no break and left for 1 h at 37°C to attach to coated wells in a humidified incubator without CO_2_ to avoid medium acidification. Seahorse XF96 plate (101085-004, Agilent) was used following the manufacturer’s instructions for XF Cell Mito Stress Test Kit (103015-100, Agilent). In this test, sequentially, 1 μM oligomycin, 2 μM FCCP and 0.5 μM rotenone / antimycin A were added to the media. Oxygen consumption rate (OCR) and extracellular acidification rate (ECAR) data were recorded by WAVE software (Agilent). OCR and ECAR data were normalized to cell count and first baseline measurement of WT cells. Basal, maximum and spare respiratory capacities were extracted with area under curve analysis in GraphPad Prism.

### 2.15 Analysis of mitochondria

For TMRE staining, B cells were washed in 150 μL PBS, stained with 1:500 Zombie Violet for dead cell discrimination in PBS on ice, washed 2 × 100μL with complete RPMI and stained with 5 nM TMRE (T669, Thermo Fisher Scientific) in 200 μL complete RPMI at RT for 20 min. Resuspended in 150 μL of complete RPMI, cells were immediately analyzed by flow cytometry, on BD LSR Fortessa.

For Tom20 staining, B cells were stained with Zombie Violet as described above, fixed with 1.6% formaldehyde in PBS for 10 min, washed 2 × 150 μL PBS, permeabilized with 0.5% Triton X-100 in PBS for 5 min at RT, blocked for 1 h at RT. Incubation with primary Tom20 antibodies was done at 1:500 dilution for 30 min, followed by 3 × 150 μL washes, staining with 1:1000 dilution of anti-rabbit-AF488 secondary antibodies, and 3 × 150 μL washes. Cells were then resuspended in 130 μL and analyzed by flow cytometry, on BD LSR Fortessa. Antibody incubations, blocking and washings were done in flow cytometry buffer I on ice. Geometric mean fluorescence intensities were extracted with FlowJo software.

### 2.16 Statistics and data presentation

Statistical analysis was performed in GraphPad Prism. Student’s t test was applied to the data comparing WT and MIM-KO groups. Antibody titres and microscopy data were analyzed with unpaired two-tailed t test unless otherwise stated. Additionally, for TIRF microscopy datasets, geometric means were extracted for each biological replicate and means were analyzed by ratio paired t test. In other experiments, ratio paired t test was also applied when pairing WT and MIM-KO data was based on the day of the experiment as indicated in figure legends. Multiple measures two-way ANOVA was additionally used to compare the antibody responses upon immunization. Data presented as Mean ± SEM, unless stated otherwise. Significance is denoted as * p<0.05, ** p< 0.01, *** p< 0.001, **** p<0.0001. Inkscape and Adobe Illustrator were used for figure assembly, and Biorender for schematics.

## 3 Results

### 3.1 Largely normal B cell development and maturation of B cells in MIM^−/−^ mice

Mice with targeted disruption of *Mtss1* gene (MIM^−/−^), lacking the expression of MIM, were generated previously (Saarikangas et al., 2011). Splenic B cells, that normally show robust MIM staining in immunoblot, isolated from these mice showed no detectable MIM expression (Fig. 1A). To investigate the possible functions of MIM in the B cell compartments, we first examined the B cells in the bone marrow. We notified a slightly increased number of hematopoietic progenitors, as defined by CD117^+^ CD19^−^ cells but found no apparent differences in the numbers of CD19^+^ and CD19^+^ IgM^+^ populations between age-matched wild type (WT) and MIM^−/−^ mice (Fig. 1B-C, Suppl. Fig. S1B). We then carried out more detailed analysis of the bone marrow B cells with additional surface markers to resolve consecutive developmental stages from common lymphoid progenitors to immature and mature recirculating B cells (gating strategy shown in Suppl. Fig. S1A). We found no significant differences between WT and MIM^−/−^ mice in any of these developmental stages (Fig. 1D).

**Figure 1.**
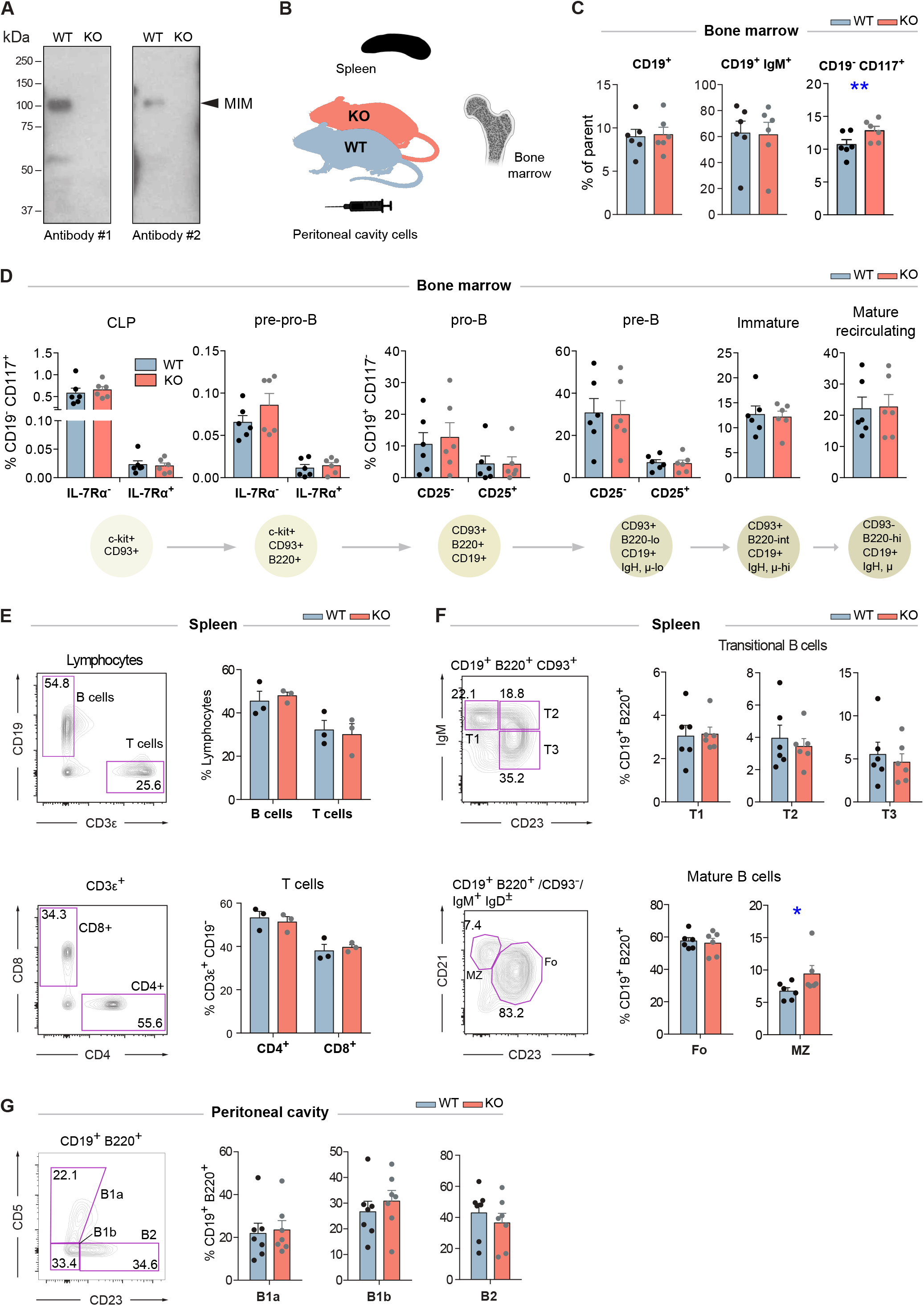
Largely normal B cell development and composition of B cell compartments in the bone marrow, spleen and peritoneal cavity of MIM^−/−^ mice. **(A)** Equal numbers of WT and MIM-KO splenic B cells were lysed and analyzed by immunoblotting with MTSS1 antibodies (4385S (Ab #1), 4386S (Ab #2), Cell Signaling Technology). Arrowhead indicates the position of the band corresponding to MIM. **(B)** Bone marrow, spleen and peritoneal cavity cell populations from WT and MIM^−/−^ mice were extracted for analysis by flow cytometry (C-G) using the gating strategy shown in Supplementary Fig. S1 and S2. Data of 3–6 independent experiments is shown as mean ± SEM. **(C)** Percentages of total CD19^+^, CD19^+^ IgM^+^ and CD19^−^ CD117^+^ cells in the bone marrow. **(D)** Percentages of B cell precursor and mature recirculating B cell populations in the bone marrow. Progression through consecutive B cell developmental stages was analyzed based on the major surface phenotypic markers shown in the schematic below. Percentage of corresponding bone marrow populations are shown as mean ± SEM. **(E)** Percentages of CD19^+^ B cells and total (upper panel) as well as CD4^+^ and CD8^+^ T cell populations (lower panel) in the spleen. (**F)** Percentages of major B cell subsets in the spleen. T1–3 (transitional 1–3), Fo (follicular), MZ (marginal zone) B cells are analyzed. (**G)** Percentages of CD23^−^CD5^+^ (B1a), CD23^−^CD5^−^ (B1b), CD23^+^CD5^−^ (B2) B cells in the peritoneal cavity.

Next, we went on to analyze the maturation of B cells and their different subsets in the periphery (gating strategy shown in Suppl. Fig. S2A, B). No defects were observed in the overall percentages of CD19^+^ B cells or major T cell subsets in the spleen (Fig. 1E). Also the proportions of transitional (T1–3) and follicular (Fo) cells were not significantly altered, while marginal zone (MZ) B cells appeared slightly elevated (Fig. 1F). A special self-renewing population of B cells concentrate in the peritoneal and pleural cavities. To analyze these B1 cells, we isolated cells from the peritoneal cavity of MIM^−/−^ and WT mice. We found no significant differences in the proportions of CD5^+^ (B1a), CD5^−^ (B1b) or mature peritoneal B cells (B2) (Fig. 1G). These results demonstrate that the development of different B cell subsets, both in the bone marrow as well as in the periphery, does not depend on MIM.

### 3.2 MIM^−/−^ B cells are defected in BCR signaling upon activation with surface-bound antigens

Previous cell biological studies in other cell types, supported by various biochemical assays, have proposed a role for MIM at the interface of the plasma membrane and the actin cytoskeleton (Lee et al., 2007; Lin et al., 2005; Mattila et al., 2003, 2007; Saarikangas et al., 2011, 2015). The actin cytoskeleton is intimately involved in the activation of B cells on antigen presenting cells by enabling the spreading of the cells and the formation of the immunological synapse (Harwood and Batista, 2010). Interestingly, the organization of the plasma membrane and the actin cytoskeleton have also been shown to regulate the signaling capacity of the BCR (Treanor et al., 2010; Mattila et al., 2013, 2016). To examine the role of MIM in BCR signaling, we, first, analyzed the mobilization of intracellular calcium, one of the most dramatic immediate consequences of BCR triggering. We isolated splenic B cells from WT and MIM^−/−^ mice, loaded them with Fluo-4 and Fura Red, and stimulated with decreasing concentrations of surrogate antigen, either full antibodies against mouse IgM BCR (anti-IgM), or their F(ab’)_2_ fragments, while analyzing the response by flow cytometry. Ratiometric analysis of Fluo-4/Fura Red fluorescence intensities revealed similar elevation of intracellular Ca^2+^ levels in both WT and MIM^−/−^ B cells (Fig. 2A, Supplementary Fig S3F). We also analyzed the calcium response in peritoneal cavity B cells, which were loaded with Fluo-4, prestained with CD23-Alexa Fluor (AF)-594 just before acquisition and stimulated with AF633-labeled anti-IgM antibodies, allowing distinction between B1 (IgM^+^ CD23^−^) and B2 (IgM^+^ CD23^+^) cells. Intracellular Ca^2+^ mobilization was found comparable between WT and MIM^−/−^ cells also in these two B cell subsets (Suppl. Fig. S3A).

**Figure 2.**
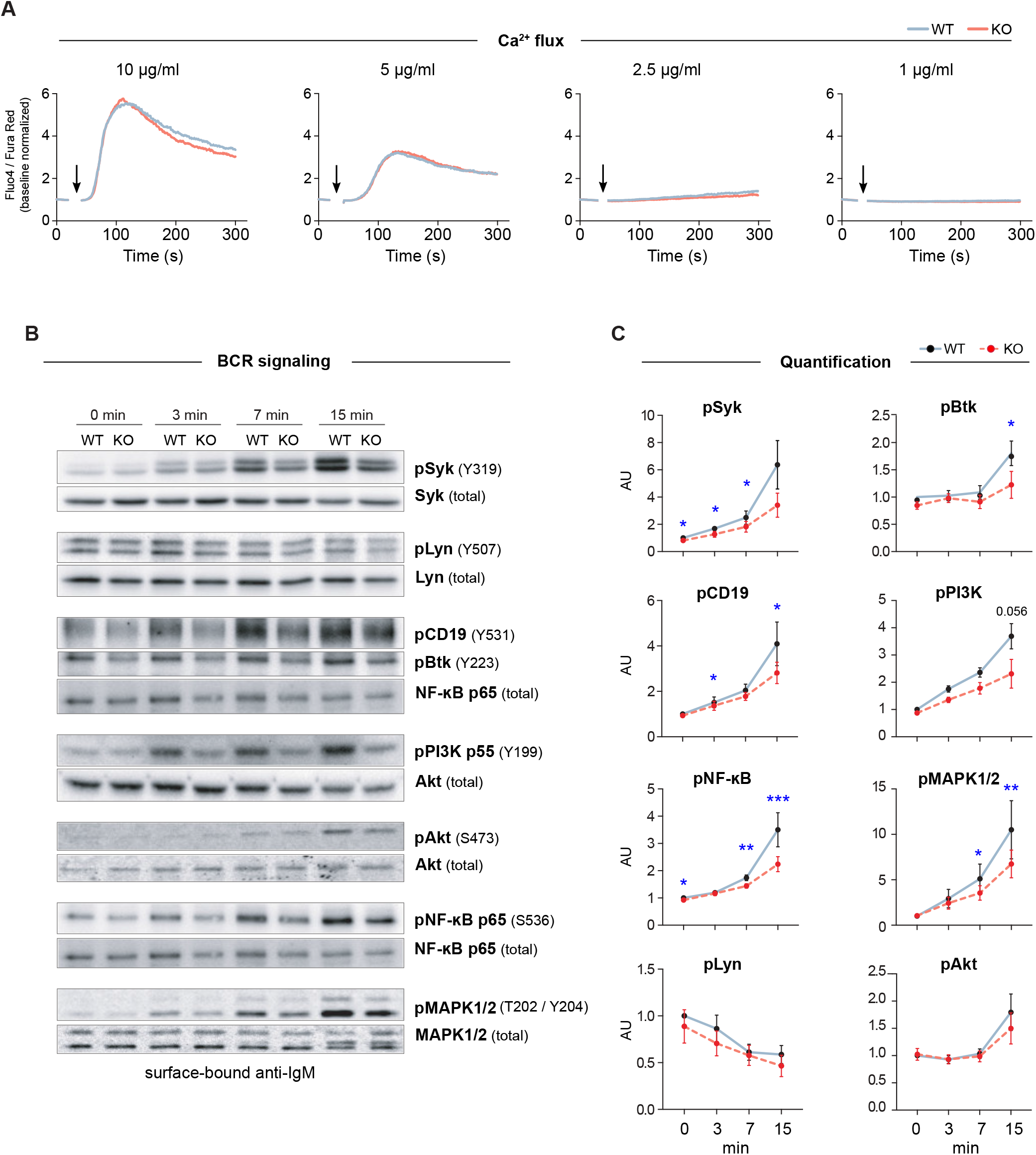
MIM deficiency leads to impaired B cell receptor signaling in response to surface-bound antigen. **(A)** Flow cytometry analysis of Ca^2+^ mobilization in response to BCR stimulation in solution. Splenic B cells from WT or MIM-KO mice were labeled with Fluo-4 and Fura Red and stimulated with 10, 5, 2.5 or 1 μg/mL anti-IgM antibodies. Time of anti-IgM antibody addition is indicated by an arrow. Data is presented as a ratio of Fluo-4 to Fura Red median fluorescence intensity. Mean values of 3-5 independent experiments are shown. **(B)** Splenic B cells were stimulated on surfaces coated with 5 μg/mL anti-IgM antibodies for 0, 3, 7 and 15 min and lysed. Lysates were subjected for immunoblotting for phosphorylated forms of different BCR signaling effector proteins. Unphosphorylated forms of the corresponding proteins were used as loading control, except for pCD19, pBtk and pPI3K, where NF-κB p65, Syk or Akt were used as loading controls. **(C)** Quantification of the data in (B). Data is presented as ratios of phosphorylated forms to total protein levels and normalized to the level of WT at 0 min. Data is from 4–8 independent experiments. Mean ± SEM is shown. * p<0.05, ** p< 0.01, *** p< 0.001.

To study the BCR signaling in more detail, we next analyzed activation of individual components of BCR signaling pathway by looking at phosphorylation levels of downstream effector molecules. Splenic B cells were stimulated with soluble or surface-bound anti-IgM antibodies for 3, 7 and 15 min and analyzed by immunoblotting. As expected, both stimulatory conditions induced rapid activation of BCR signaling components, as detected by the levels of phosphorylated signaling proteins. Importantly, MIM^−/−^ B cells showed clear defects in signaling in response to surface-bound anti-IgM. Most of the analyzed molecules, including Syk, CD19, Btk, p65 NF-κB and MAPK1/2 showed significant reduction in their activation (Fig 2B, C). Consistently, we found similar differences when using either 5 or 1 μg/ml of surface-tethered anti-IgM, or corresponding amounts of F(ab’)_2_ fractions of anti-IgM (Suppl. Fig. S3D-E). The signaling components studied can be classified into different cascades from the proximal players to downstream effectors. While MIM^−/−^ cells showed robust defects in proximal signaling molecules, the extent to which they affected the downstream cascades alternated. The defects in PI3K pathway were largely recovered at the level of Akt. At the same time, the levels of pp65 NF-κB and pMAPK1/2 were decreased, suggesting that MIM^−/−^ B cells were inefficient in triggering the diacylglycerol (DAG)-PKC module, targets of which both NF-κB and MAPK1/2 are (Mérida et al., 2010). Interestingly, when we studied activation of BCR by soluble ligand, anti-IgM surrogate antibodies in solution, MIM^−/−^ B cells only showed significant defects in the activation of the proximal kinase Syk, but normal activation of other signaling components (Suppl. Fig S3B, C). These results place MIM as a regulator of, specifically, BCR signaling by surface-bound antigen, function that clearly depends on the fine-tuned activities of the actin cytoskeleton (Bolger-Munro et al., 2019) and could fit well with the previously postulated role of MIM as an organizer of the actin cytoskeleton-membrane interface.

### 3.3 The morphology and formation of the immunological synapse is unaltered in MIM^−/−^ B cells

The actin cytoskeleton is one of the major organizers of cell shape. To explore whether MIM^−/−^ B cells showed any changes in the overall morphology, we visualized them using scanning electron microscopy (SEM), either in resting state or after 10 min activation by surface-tethered anti-IgM, mimicking the formation of the immunological synapse. In SEM, the morphology of MIM^−/−^ B cells appeared grossly similar to WT cells (Fig. 3A). However, to perform another, more quantitative analysis, we turned to fluorescent microscopy. We activated B cells on anti-IgM-coated coverslips and analyzed the area of spreading using total internal reflection fluorescence (TIRF) microscopy, an imaging method ideal for visualization of the cell membrane-coverslip interface. The analysis of the TIRF images revealed that MIM^−/−^ B cells spread slightly but consistently less than their WT counterparts (Fig. 3B). To measure overall phosphorylation at the contact region, we stained the cells with anti-phospho-Tyrosine (pTyr) antibodies. Mean fluorescence intensities of pTyr were also reduced in MIM^−/−^ B cells. To investigate the impaired BCR signaling and its kinetics in more details, we next activated the B cells on anti-IgM coated coverslips for different timepoints and stained them with anti-pBtk and anti-pSyk antibodies, as well as phalloidin. Quantification of the pBtk and pSyk in the contact plane showed significantly reduced levels of these signals, as well as levels of spreading, in MIM^−/−^ cells through different time points (Fig 3C). The diminished area of spreading and kinase signaling detected by microscopy are well in line with our results from immunoblotting that also showed impaired BCR signaling. However, based on the SEM data and F-actin staining of the splenocytes, MIM^−/−^ cells do not exhibit major morphological defects in their actin cytoskeleton (Fig. 3A–C).

**Figure 3.**
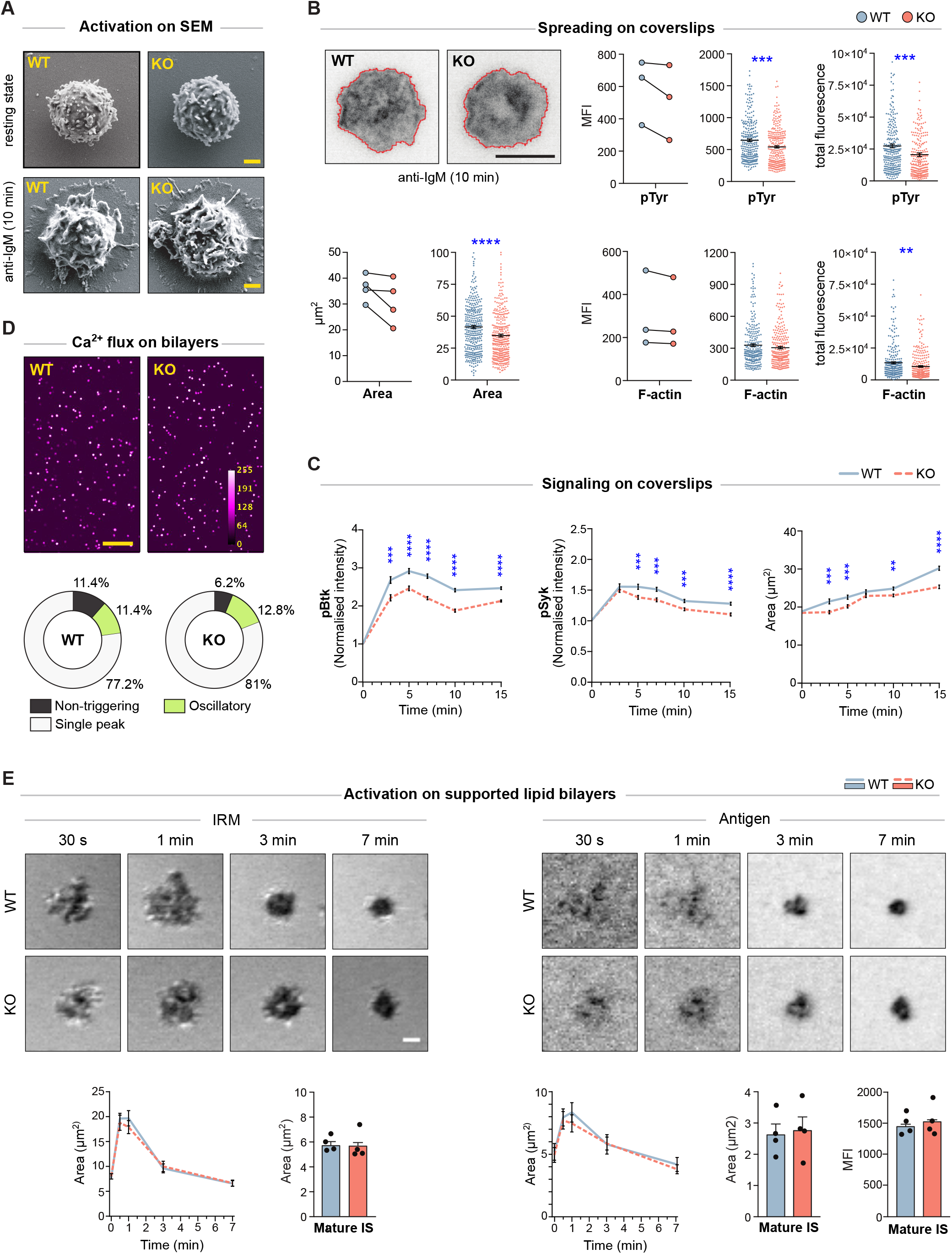
MIM^−/−^ B cells show diminished spreading and signaling on antigen-coated glass, but gather normal levels of antigen on supported lipid bilayers. **(A)** Scanning electron micrographs of WT and MIM-KO splenic B cells at resting state, let to adhere on CellTak-coated coverslips, or let to spread and get activated on coverslips coated with 5 μg/mL anti-IgM for 10 min. Scale bar, 1 μm. **(B)** Splenic B cells were stimulated on coverslips coated with 7.5 μg/mL anti-IgM antibodies for 10 min, fixed, permeabilized and stained with phalloidin (F-actin) and anti-phospho-Tyrosine antibodies (pTyr). Samples were imaged with TIRF microscopy. Representative images showing cell spreading and F-actin staining, are shown (upper left). Area of spreading was analyzed by signal thresholding in pTyr channel (lower left panels), and pTyr (upper right panels) and F-actin (lower right panels) intensities inside cell perimeter were quantified using ImageJ. Mean (MFI) and total (total) fluorescence intensities of pTyr and F-actin stainings are shown, and the mean fluorescence intensity is presented both as pairwise comparison of geometric means of individual experiments (on the left) and as scatter plots of random sampling of 92 cells from each of 3-4 individual experiments (middle). Mean ± SEM is shown. Scale bar, 5 μm. **(C)** Splenic B cells were stimulated on coverslips coated with 5 μg/mL anti-IgM antibodies for 3, 5, 7, 10 and 15 min, fixed, permeabilized and stained with phalloidin (F-actin) and anti-phospho-Btk (pBtk) or anti-phospho-Syk (pSyk) antibodies. Samples were imaged with spinning disk confocal microscopy. Area of spreading was analyzed by signal thresholding in phalloidin channel, and pBtk or pSyk mean fluorescence intensities (MFI) inside cell perimeter were quantified using ImageJ and normalized to non-activated cells (time 0 min). Normalized MFI of pBtk and pSyk, and spreading area based on F-actin are shown. Data is shown as mean ± SEM of 4 independent experiments (~25-150 cells per condition per experiment). WT and KO cells were compared using unpaired t-test. **(D)** Intracellular Ca^2+^-flux was analyzed in splenic WT and MIM-KO B cells loaded with Fluo-4 and stimulated with anti-kappa light chain antibodies tethered on supported lipid bilayers (SLB). A spinning disk confocal microscope was used to record the Fluo-4 intensity and the intracellular Ca^2+^ levels were quantified with *CalQuo^2^* software. Representative images of stimulated B cells, 85 sec after injection into the chamber are shown (upper panel). Mean percentages of non-triggering cells and cells with single peak or oscillatory responses are shown (lower panel). Data of 3 independent experiments. Scale bar, 100 μm. **(E)** Splenic WT and MIM-KO B cells were stimulated with monobiotinylated anti-kappa light chain antibodies tethered via AF647-labelled streptavidin on SLBs. A laser scanning confocal microscope was used to detect IRM signal (panels on the left) corresponding to the area of contact and to measure the amount of antigen collected in the synapse from the AF647 signal (panels on the right). The data was quantified for cell area (graphs on the left), antigen area and antigen fluorescence intensity (graphs on the right) during synapse formation (0-7 min) by tracking individual cells from live cell imaging movies, and in the mature synapse (>10 min) by taking snapshots from several fields of views after movie acquisition. Data is from 3-4 independent experiments (~25-30 cells per condition in the movie, ~400-450 cells in the snapshots). Data were compared using unpaired t-test. Mean ± SEM is shown. Scale bar, 1 μm. * p<0.05, ** p< 0.01, *** p< 0.001, **** p<0.0001

Recognition of surface-bound antigens *in vivo* can involve interaction of B cells with antigenic determinants on substrates of various physical properties. While stiff substrates can include bacterial cell wall components or extracellular matrix-linked antigens, perhaps more typical encounter occurs on the surface of antigen presenting cells, where antigens remain laterally mobile. To understand if MIM^−/−^ B cells can initiate robust BCR signaling upon encounter with mobile antigens, we first settled Fluo-4-loaded WT and MIM^−/−^ B cells on AF555-labeled supported lipid bilayers (SLB), containing anti-kappa antibodies and ICAM-1. Mobilization of Ca^2+^ was imaged for 5 min by spinning disk confocal microscopy and analyzed with the *CalQuo^2^* software package for MATLAB (Lee et al., 2017). Analysis of median Fluo-4 intensity and proportions of MIM^−/−^ B cells with single peak or oscillatory Ca^2+^ intensity profiles revealed no differences between MIM-KO and WT cells (Fig. 3D). Next, we evaluated the ability of MIM-KO cells to spread and form immunological synapses on SLBs. We performed live-cell imaging of WT and MIM-KO cells and recorded interference reflection microscopy (IRM) images to detect the overall contact area of the cells, and fluorescent antigen signal to follow antigen gathering in the synapse (Fig. 3E). The movies showed fast cell spreading, reaching maximum in 1-2 minutes, followed by slower contraction response coinciding with antigen accumulation to form the mature immunological synapse. We did not detect any differences in the kinetics of synapse formation in MIM-KO cells as compared to WT counterparts. Also, in the mature synapses, analyzed after 10 minutes of contact with the SLB, MIM^−/−^ B cells accumulated normal amounts of antigen in the center of the synapse (Fig. 3E). Thus, while MIM modulates BCR signaling and spreading responses upon antigen contact on stiff substrates, upon engagement of membrane-bound mobile antigens the cells are able to engage and gather normal amounts on antigen to the center of the immunological synapse. On membranous surfaces, spreading and contraction response is complemented with dynamic fluctuations of F-actin-rich protrusions, making the regulation of the cytoskeletal structures distinct from spreading on immobilized antigens.

### 3.4 MIM is required for an efficient antibody response against T-independent antigen

To test if the defected signaling in MIM^−/−^ B cells leads into problems in mounting of the immune responses, we went on to examine antibody levels of our mice first at the basal state and then upon immunization with T cell-independent (TI) or T cell-dependent (TD) model antigens. We saw no significant changes in the basal antibody levels in the sera of WT and MIM^−/−^ mice (Fig. 4A). To study the development of antibody responses towards TI antigens, we immunized mice with NP-FICOLL (Fig. 4B–D). Throughout the examination time course of 28 days, we found reduced levels of both total and NP-specific IgM in MIM^−/−^ mice (Fig. 4C). Interestingly, we also detected impaired responses in the total levels of IgG subtypes, most profound in IgG2b, while the production of NP-specific IgG subclasses was not impaired (Fig. 4D).

**Figure 4.**
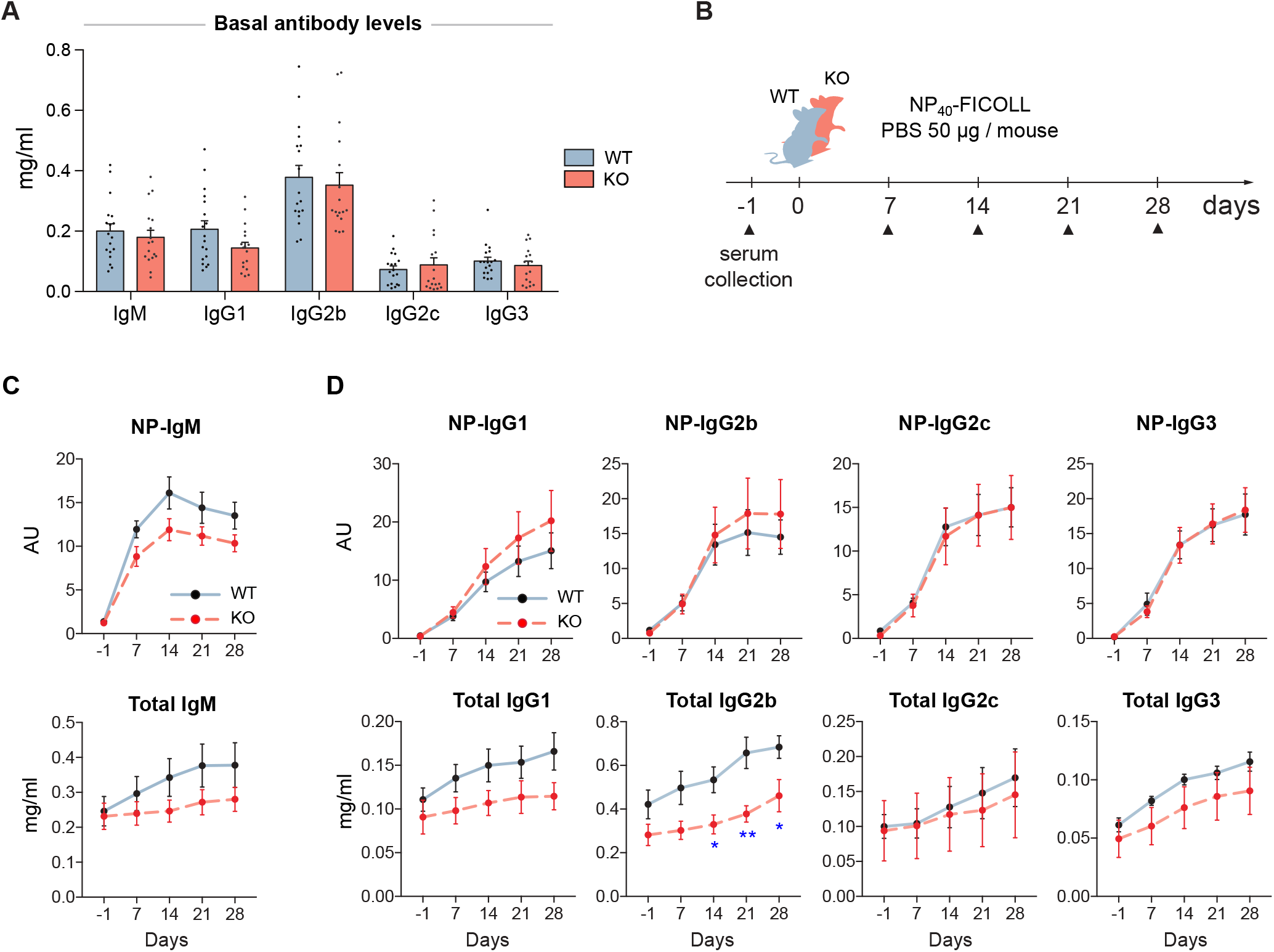
MIM-deficiency results in impaired IgM response and reduced levels of total IgGs during T cell-independent immune response. **(A)** Basal antibody levels of major immunoglobulin subclasses were measured from 17–18 WT and MIM KO mice. **(B)** A schematic representation of T cell-independent (TI) immunization study. **(C)** The antibody responses of immunized WT and MIM-KO mice were followed from serum samples as in (B). Total and NP-specific antibody levels of different immunoglobulin isotypes were measured with ELISA. 7-8 mice per group. Mean ± SEM is shown. * p<0.05, ** p< 0.01.

For immunizations with TD protein antigen, we used NP-KLH in alum and followed up the primary response for four weeks. Four months later, we stimulated a memory response and followed that for 2 weeks (Suppl. Fig. S4A). In contrast to TI immunization scheme, we found that MIM^−/−^ mice were as efficient in mounting antibody responses to a protein antigen as their WT counterparts (Suppl. Fig. S4B-C). Analysis of the memory responses also showed equal levels of NP-specific IgG for all subclasses (Suppl. Fig. S3D). Furthermore, we analyzed affinity maturation by comparing binding to low and high densities of NP-epitopes in ELISA, and found no significant differences between WT and MIM^−/−^ mice (Suppl. Fig. S4E).

In the light of defects in BCR signaling and TI immune responses in MIM^−/−^ mice, normal antigen-specific IgG immune responses may point towards compensation by other signals in the system, such as T cell help. To dissect the B cell intrinsic features linked to IgG antibody responses in more detail, we set up an *in vitro* assay for class-switch recombination (CSR). We provoked B cells to change the isotype of the produced Ig molecules, by mimicking cellular events of pathogen encounter or T cell help during maturation of IgG antibody responses. As expected, 3 days of B cells activation with LPS or CD40L in combination with cytokines induced switching of surface-expressed Ig molecules to different IgG isotypes, as detected by flow cytometry. Consistent with the *in vivo* data on generation of NP-specific IgG antibody responses, MIM^−/−^ B cells switched normally in all tested conditions, producing similar percentages of IgG^+^ cells, indicating that MIM is not required for CSR in response to TLR ligands or CD40L and cytokines. In fact, switching rates into the most common isotypes were even slightly higher for MIM^−/−^ B cells when stimulated with LPS (Suppl. Fig. S5A–D). We also loaded the cells with Cell Trace Violet dye, which allowed us to analyze fluorescence profiles of dividing splenic B cells in response to these stimuli. We found robust proliferation of MIM^−/−^ B cells when they were activated with LPS, CD40L or anti-IgM + CD40L (Suppl. Fig. S5E). Analysis of the proliferation indices (PI) showed close to equal proliferation of MIM-KO and WT cells in all conditions tested. Notably, the division indices (DI) of MIM^−/−^ B cells were moderately, but significantly, increased in CD40L + IFNg cultures, reflecting smaller numbers of cells left undivided, further indicating normal or even improved ability of MIM^−/−^ cells to react to conditions similar to T cell help.

Our immunization studies suggest that MIM plays a role in IgM antibody responses to TI type 2 antigens, which haptenated FICOLL represents. However, MIM is dispensable for the development of antibodies against TD protein antigens. The ability to switch the antibody class upon polyclonal activation with LPS, TI type 1 antigen, appeared normal. Although by using a conventional MIM knockout mouse model we cannot rule out the influence of other immune and stromal cells on the immune responses generated *in vivo*, IgM antibody responses to TI antigens are considered to be largely B cell intrinsic and in case of TI type 2 antigens rely primarily on intact BCR signaling. Thus, the defects in IgM antibody responses are in line with the diminished BCR signaling observed *in vitro* (Fig. 2B–C; Fig 3B–C).

### 3.5 MIM^−/−^ B cells show higher metabolic profile upon LPS and CpG stimulation

Our results suggest that MIM plays a role in antibody responses to TI-2 antigens. *In vivo*, however, T cell-independent antigens are usually associated with pathogenic determinants recognized by Toll-like receptors, important modulators of B cell immune responses (Bekeredjian-Ding and Jego, 2009). Activation of resting lymphocytes upon BCR engagement or recognition of microbial components triggers dramatic changes in their metabolism. Upregulated metabolic activity meets the increased energetic demands of activated cells and supports active proliferation and differentiation into antibody-secreting cells. We next analyzed the metabolic reprogramming potential of MIM^−/−^ B cells upon activation via either BCR or TLRs, or combination of both. The rates of cellular oxygen consumption (oxygen consumption rate, OCR), an estimate of mitochondrial respiration, and extracellular acidification (extracellular acidification rate, ECAR), an estimate of glycolysis, were measured with a Mito Stress test in the Seahorse platform. In this assay, serial injections of oligomycin, Carbonyl cyanide-4-(trifluoromethoxy)phenylhydrazone (FCCP) and rotenone/antimycin A (Rot/AA) mixture, sequentially inhibit ATP production, collapse the proton gradient membrane potential and finally shut down mitochondrial respiration. This system allows measurement of the basal OCR, maximal and spare respiratory capacities, as well as determination of non-mitochondrial respiration.

First, we analyzed metabolic responses to individual stimulations with BCR ligands or TLR agonists. To this end, we activated WT and MIM^−/−^ B cells with LPS, CpG or anti-IgM + IL-4 for 24 h or kept them largely resting with IL-4 alone (Fig. 5A). Already samples treated with IL-4 alone showed a slight but significant increase in the maximal respiration in MIM^−/−^ B cells. Notably, we observed a significant increase in the metabolic profile of MIM^−/−^ cells as compared to WT counterparts upon activation with TLR4 and TLR9 ligands, LPS or CpG. Approximately 20% increase in the maximum respiration of MIM^−/−^ cells was measured upon treatment with LPS or CpG. MIM^−/−^ cells also showed similar increase in the basal OCR levels and almost two-fold upregulation for spare respiratory capacity upon CpG activation. In contrast to TLR stimulations, 24 h activation via BCR led to a slight diminution in basal respiration and no differences in maximum and spare respiration levels in MIM^−/−^ cells. The preferred route of ATP generation, reflected by basal OCR to ECAR ratio, remained relatively constant across conditions, indicating no major shifts in the balance of oxidative phosphorylation and glycolytic metabolism.

**Figure 5.**
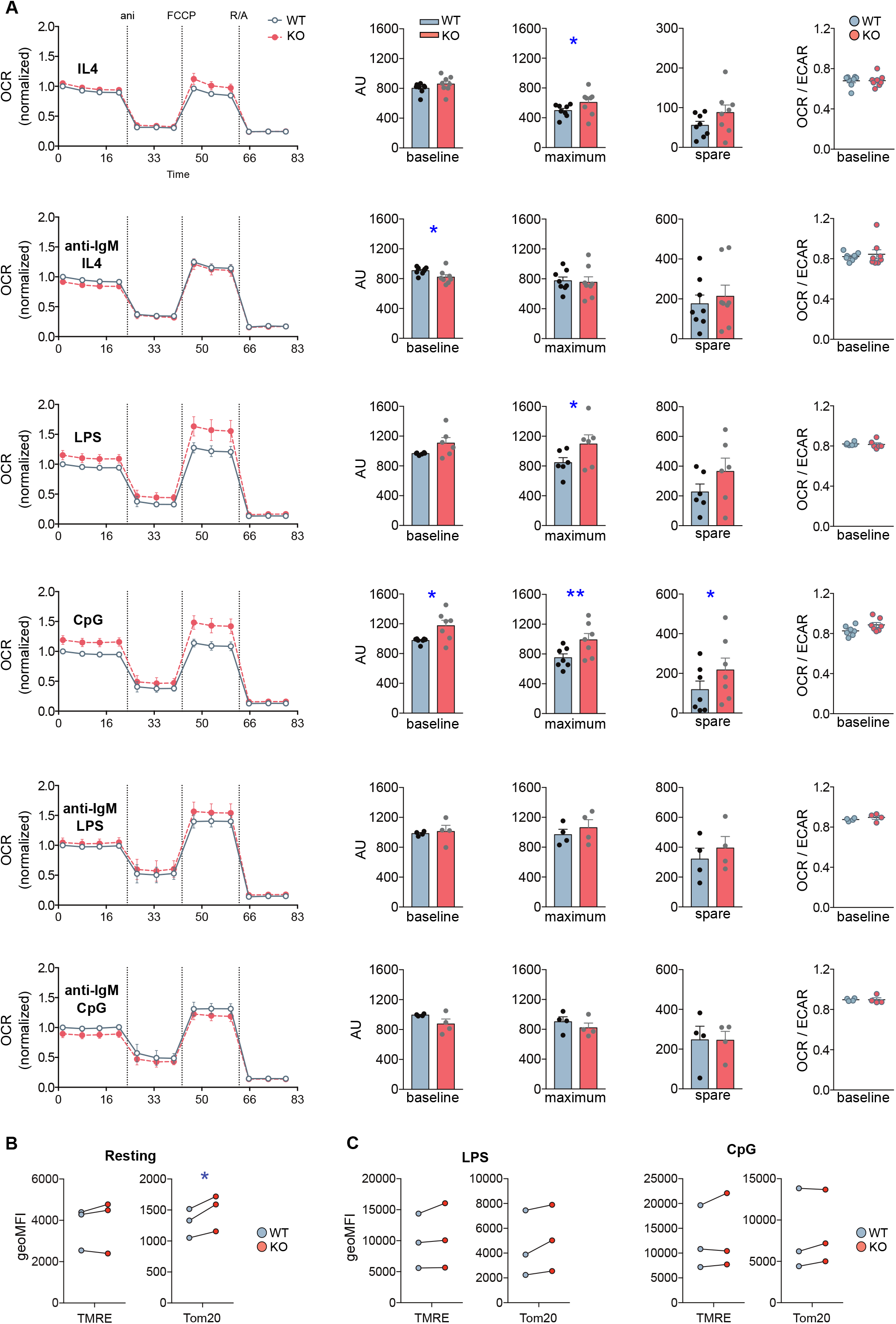
MIM-deficient B cells show increased metabolic activity upon stimulation with Toll-like receptor ligands LPS and CpG. **(A)** Oxygen consumption rate (OCR) profiles of WT and MIM-KO splenic B cells stimulated with IL-4, anti-IgM + IL-4, LPS, CpG, anti-IgM + LPS, or anti-IgM + CpG for 24 h were measured in a Seahorse XF Cell Mito Stress Test assay. The different steps of the assay are depicted in the graph on the left showing mean ± SEM. Comparisons of baseline mitochondrial respiration as well as maximum and spare respiratory capacities, extracted from the assay, are shown in the graphs in the middle, and quantification of the ratio of OCR to extracellular acidification rate (ECAR) at the baseline is shown on the right. Data presented as mean of 4–8 independent experiments. Mean ± SEM. (**B–C)** Splenic B cells, either left unstimulated (**B**) or stimulated with LPS or CpG for 24 h (**C**), were loaded with TMRE to probe for the mitochondrial membrane potential, or stained with anti-Tom20 antibodies to derive total mitochondrial mass. The samples were analyzed by flow cytometry. Data of 3 independent experiments. Mean ± SEM. * p<0.05.

To gain more understanding into the relationship of BCR and TLR signals in the activation of MIM^−/−^ cells, we then asked how the metabolic reprogramming is affected by simultaneous engagement of BCR and TLR. To this end, we cultured WT and MIM^−/−^ B cells for 24 hrs with combination of anti-IgM + LPS or anti-IgM + CpG and analyzed their metabolic profiles (Fig. 5A). Surprisingly, we found that TLR-mediated increase in oxidative metabolism in MIM^−/−^ cells was restored to WT levels by simultaneous BCR engagement. To more directly compare this data to anti-IgM-stimulated cells, where IL-4 was added to increase survival of the cells, we carried out analysis of anti-IgM + LPS/CpG - stimulated cells also in the presence of IL-4. We found that simultaneous engagement of BCR also in these conditions reverted the TLR-mediated increase in oxidative metabolism in MIM^−/−^ B cells closer to WT levels (Suppl. Fig. S6A). However, the negative effect of anti-IgM was less efficient in presence of IL-4, consistent with the finding that IL-4 supports metabolic reprograming towads higher oxidative state levels in MIM^−/−^ cells.

Finally, we asked if the higher metabolic activity of MIM^−/−^ B cells reflects a *bona fide* change in the mitochondrial activity or is a result of elevated mitochondrial biogenesis upon TLR-mediated metabolic reprograming. To this end, we stained mitochondria with TMRE and antibodies against Tom20 to assess mitochondrial membrane potential and mitochondrial mass, respectively. In freshly isolated splenic MIM^−/−^ B cells, we found slightly but significantly elevated mitochondrial mass without significant increase in mitochondrial membrane potential (Fig. 5B). As expected, activation of B cells with either anti-IgM + IL-4, LPS or CpG increased mitochondrial biogenesis and membrane potential, with CpG inducing most potent response (Fig. 5C). However, WT and MIM^−/−^ B cells showed comparable levels of mitochondrial membrane potential as well as mitochondrial mass after TLR stimulation. This suggests that the changes we observed in the metabolic activity of MIM^−/−^ cells occur without detectable additional mitochondrial biogenesis but are rather a function of increased efficiency that might not be adequately represented by TMRE. Functional changes in the mitochondrial, and metabolic, activity may also be reflected by cell size (Miettinen and Björklund, 2017). We found that CpG-but not LPS- or anti-IgM + IL-4-activated MIM^−/−^ cells often showed slightly increased forward scatter characteristics when analyzed by flow cytometry (Suppl. Fig. S6B). Together, our results suggest that in response to TLR4, TLR9 or IL-4 stimulation, MIM negatively regulates metabolic reprograming of B cells, decreasing reserves of oxidative metabolism, which are accompanied by functional changes in the mitochondrial activity, rather than mitochondrial biogenesis. Interestingly, in MIM^−/−^ B cells, simultaneous engagement of BCR reverts TLR-mediated increase in oxidative mitochondrial activity.

## 4 Discussion

In this study, we examined the enigmatic cytoskeleton-membrane linker protein MIM in B cells, where it is strongly expressed and also implicated in B cell lymphomas (Yu et al., 2012). In our mouse model, we found diminished TI antibody responses, as well as defected BCR signaling, indicative of a role in B cell activation and mounting of humoral immune responses. Together with the upregulated metabolic boost and normal proliferative responses in MIM^−/−^ B cells upon TLR activation, our data suggest a complex role for MIM in modulation of different signaling pathways and cell fitness.

The predisposition of another MIM knockout mouse strain to lymphomagenesis, reported by Zhan and colleagues, is intriguing (Yu et al., 2012). We found the spleens of MIM^−/−^ mice at the age of 2-7 months normal in size and cellularity, but as we did not perform aging experiments, we cannot preclude problems later in life. Regarding B cell development, Zhan and colleagues at first showed aberrant levels of total CD19^+^ or CD19^+^ IgM^+^ cells in lymphoid organs (Yu et al., 2012), yet later reported normal B cell numbers (Zhan et al., 2016). Following the same gating strategy, the numbers of pre-B cells in our MIM^−/−^ strain appeared normal (Suppl. Fig. S1B–C), and we postulate that the observed discrepancies may arise from differences in strain maintenance or mouse age at the time of immunophenotyping. The possible consequences of the deletion of MIM for B cell development might also be too subtle to manifest in all models. In our model, we however, noted mildly elevated numbers of MZ B cells in the spleens of MIM^−/−^ mice (Fig. 1F). The mouse strain generated by Zhan and colleagues used embryonic stem (ES) cells with insertion of gene trap sequence between exons 3 and 4 (clone CSC156, BayGenomics) (Yu et al., 2011). Notably, an independent attempt to recapitulate the generation of MIM^−/−^ strain with the same clone of ES cells resulted in an inefficient use of delivered splicing acceptor site (Fahrenkamp et al., 2017). This left a considerable expression of full-length Mtss1 both at mRNA and protein levels, and varible remaining expression levels between animals, indicative of lability of the splicing. Lack of MIM expression in analyzed cells was, however, shown by Zhan and colleagues (Yu et al., 2011, 2012). In our strain, where a neomycin cassette with several stop codons is inserted in exon 1 of *Mtss1*, low levels of alternative splicing have been previously observed. However, the generated transcript translates into a N-terminally truncated polypeptide chain, where the critical I-BAR domain is functionally inhibited, and which did not yield detectable levels of protein expression (Saarikangas et al., 2011). In this study, confirmative of the lack of MIM expression in B cells, we also confirmed a clear and repeatable loss of detectable protein expression in our substrain, using two different commercial antibodies (Fig. 1A).

MIM^−/−^ B cells exhibited consistently diminished phosphorylation of several BCR effector molecules upon stimulation of BCR with surface-bound antigen (Fig. 2B, C). This defect was in line with smaller area of spreading and reduced overall tyrosine phosphorylation detected by microscopy at the site of contact with antigen (Fig. 3B–C). Although lower amount of engaged antigen due to inefficient actin-dependent spreading response may partially contribute to the signaling defects observed in MIM^−/−^ B cells, defected BCR signaling also leads to diminished spreading, generating a feedback loop. Signaling upon soluble antigen stimulation, on the other hand, was mostly normal showing only slight diminution in the levels of pSyk (Suppl. Fig. S3B–C). This suggests that MIM participates in the ability of B cells to discriminate between different types of antigens by playing a specific role in B cell activation on surfaces. Although the differential responses of B cells to different forms of antigen are nowadays widely accepted (Mattila et al., 2016; Snapper, 2018; Bolger-Munro et al., 2019), to our knowledge there are only few molecules, such as CD19 and CD81 (Depoil et al., 2008; Mattila et al., 2013), that have been reported to specifically regulate stimulation by surface-linked antigens with mechanisms likely separate from structural roles in cell adhesion or spreading. We also found that while stimulation with surface-bound antigen resulted to a certain degree in reduced phosphorylation of all the tested BCR effectors, the defects in proximal BCR signaling, exemplified by reduced pSyk and pCD19, did not propagate evenly downstream. Reduced pCD19 seemed to largely spare PI3K pathway (Otero et al., 2001) as pAkt levels are on par with those of WT (Fig. 2B, C). At the same time, levels of both pNF-κB and pMAPK1/2 were significantly reduced, suggesting defected DAG-PKC signaling module (Su et al., 2002; Coughlin et al., 2005; Mérida et al., 2010). PKC-FAK axis has been implicated in the regulation of force-dependent B cell activation (Shaheen et al., 2017), and could be involved in the specific defects in response to surface-bound antigens that we observed. Regarding the previously suggested role for MIM in DLBCL (Yu et al., 2012), it is worth noting that dysregulated BCR signaling also plays a role in lymphomagenesis (Young and Staudt, 2013).

Our experiments on supported lipid bilayers (SLB) suggested that MIM^−/−^ B cells are able to form signaling-competent BCR-antigen microclusters and gather normal amounts of antigen in the center of the immunological synapse (Fig. 3E). Therefore, in this model system, coupling of BCR to actin and microtubule cytoskeleton, required for the cell spreading and antigen gathering, respectively, is not notably defected (Schnyder et al., 2011; Liu et al., 2012). However, results from the laterally fluid SLBs, cannot be directly compared to B cell stimulation by immobilized antigens. SLBs are also not fully comparable to situations where the cells need to overcome frictional coupling of antigen-presenting molecules to the membrane skeleton of the APC (Ketchum et al., 2014; Luxembourg et al., 1998; Dillard et al., 2014). In addition, SLBs in our experiments were functionalized with ICAM-1, which is known to lower the threshold for B cell activation (Carrasco et al., 2004).

MIM^−/−^ mice developed impaired IgM antibody responses to NP-FICOLL, but normal responses to NP-KLH immunization, indicating that the defects were specific to the nature of the encountered antigen. FICOLL polysaccharide haptenated with NP is a typical TI type 2 antigen, to which the response is thought to rely mainly on marginal zone B cells and peritoneal cavity B1b cells (Guinamard et al., 2000; Girkontaite et al., 2001; Hsu et al., 2006; Haas, 2011). As MIM^−/−^ mice showed normal proportions of peritoneal and even elevated MZ B cell compartments (Fig. 1F, G), inadequate numbers of these B cell populations are unlikely to be responsible for the reduced anti-NP IgM levels (Fig, 4C). However, further studies will be required to assess the functionality of B1 and MZ B cells in more detail. We consider the defects in possible indirect T cell help unlikely, due to the low expression of MIM in T cells (Yu et al., 2012), normal CD4^+^ T cell numbers (Fig. 1E), intact antibody responses to TD antigen NP-KLH (Suppl. Fig. S4) as well as comparable levels of class-switched IgG antibodies in response to NP-FICOLL immunization (Suppl. Fig. S5). We suggest that the impaired NP-FICOLL responses in MIM^−/−^ mice are caused by defected B cell receptor-mediated signaling, induced by cell surface-associated FICOLL molecules. This would be in line with the prevailing idea that *in vivo* B cells recognize and respond to antigens that are immobilized on the surface of other cells in the secondary lymphoid organs (Carrasco and Batista, 2007). In case of NP-KLH stimulation these defects could be rescued by second signals, to which, based on our data, MIM^−/−^ cells respond well (Fig 5, Suppl. Fig. S5, S6).

While naive, resting B cells have a considerably low metabolic profile, they elevate their metabolic rates upon activation, which typically manifested by an increase in oxygen consumption and glycolysis (Caro-Maldonado et al., 2014; Akkaya et al., 2018; Jellusova, 2018; Price et al., 2018). The mitogenic stimuli IgM, LPS and CpG have been shown to dramatically increase metabolic requirements in B cells, playing a key role in this transition, essential for differentiation into antibody secreting cells and productive immune responses (Boothby and Rickert, 2017; Jellusova, 2018).

Our results suggest that MIM plays a negative role in metabolic reprogramming of B cells towards oxidative metabolic activity upon TLR4 and especially TLR9 engagement as we found that MIM^−/−^ B cells exhibited ~20% increased metabolic activity after stimulation with either LPS or CpG, respective agonists for TLR4 and TLR9 (Fig. 5). In contrast, basal OCR levels upon activation with anti-IgM + IL-4 were reduced in MIM^−/−^ cells reflecting impaired BCR signaling observed in these cells. It is possible that STAT6-signaling triggered by IL-4 present in culture to increase the survival of the cells dilutes the effects of defected anti-IgM signaling in MIM^−/−^ cells, and leads to underestimate of the difference, as IL-4 alone lead to increased metabolic activity in MIM^−/−^ cells.

How exactly the impaired BCR signaling and abnormal metabolic rewiring of MIM^−/−^ B cells in response to TLR engagement can be explained together is an intriguing question. Surprisingly, addition of BCR engagement at the same time with TLR agonists was found to negate the metabolic phenotype found in MIM^−/−^ B cells treated with TLR agonists alone. This suggests that increased metabolic capacity in MIM^−/−^ cells could compensate the effects of compromised BCR signaling upon *in vivo* B cell activation and differentiation. However, how the different combinations of ligands are sensed by both BCR and TLRs presents complex and largely unresolved issues. In addition to distinct modes of BCR signaling induced by soluble or surface-tethered ligands, responses to simultaneous but unlinked BCR + TLR stimulations are qualitatively different from co-engagement of BCR and TLR by cross-linked ligands (Minguet et al., 2008; Sindhava et al., 2017). Thus, the exact mechanism of how MIM affects oxidative metabolic activity in response to TLR-mediated reprograming remains to be further investigated. There is, however, a large body of evidence showing that in B cells, TLR4/9-mediated responses are dependent on BCR signaling components, including BCR itself (Dye et al., 2007; Minguet et al., 2008; Jabara et al., 2012; Otipoby et al., 2015; Morbach et al., 2016; Schweighoffer et al., 2017; Sindhava et al., 2017; Suthers and Sarantopoulos, 2017). Yet, although MIM^−/−^ cells were defected in activation of CD19, PI3K, Syk, Btk, NF-κB and MAPK1/2 phosphorylation upon antigenic activation, we observed no abnormalities in proliferation of MIM^−/−^ B cells in response to either LPS or CpG (Suppl. Fig. S5E and data not shown). In the case of CpG, defects in Dock8-Src-Syk-STAT3 (Jabara et al., 2012) and PI3K/Akt-GSK3b-Foxo1 (Otipoby et al., 2015) axes primarily manifest as problems in proliferation. Inhibition of these modules, as well as MyD88-Pyk2-Lyn and CD19-PI3K-Akt (Morbach et al., 2016), in CpG-activated cells, however, leaves activation of NF-κB and MAPK1/2 largely intact. Thus, NF-κB and MAPK1/2 possess, at least partially, independent roles in TLR9 activation and may be responsible for the metabolic changes seen in CpG-activated MIM^−/−^ B cells. Indeed, NF-κB has been shown to reduce mitochondrial oxidative metabolism in sarcoma cells and MAPK1/2 have been shown to promote glycolytic shift in some cancer types, as well as in rapidly proliferating cells (Londhe et al., 2018; Papa et al., 2019). This suggests that impaired activation of these modules may underpin the elevated oxidative metabolism in MIM^−/−^ B cells. Supporting the complex relationship of metabolism with BCR and other activatory signals, it has been found that while BCR signaling initially leads to enhanced mitochondrial function, this response also rapidly shifts towards mitochondrial dysfunction if not rescued with second signals such as TLR9 agonists (Akkaya et al., 2018). Akkaya and colleagues did not study the effects of IL-4 in this context, but it seems plausible that similar rescue of the BCR-induced mitochondrial dysfunction also takes places upon IL-4 treatment. Interestingly, it has also been found, that although many B cell lymphomas depend on intact tonic or antigen-driven BCR signaling, a subset of diffuse large B cell lymphomas (DLBCL), called OxPhos DLBCL, with gene expression signature enriched in oxidative phosphorylation genes, does not express functional BCR (Caro et al., 2012; Ricci and Chiche, 2018) pointing towards separation of these signal pathways. It remains to be seen if differential expression of MIM in different cancers, including lymphomas, may correlate with changes in the metabolic activity of the cancer cells.

## 5 Acknowledgements

We are thankful for Laura Grönfors for technical assistance, Elmeri Kiviluoto for assistance in image analysis, Dr Maria Georgiadou and Prof Johanna Ivaska for their help and generosity with Seahorse assays, as well as Dr Diana Toivola and Joel Nyström for their help and generosity regarding the reagents for studying mitochondria. Central Animal Laboratory of University of Turku is acknowledged for good care of the mice. We thank Tero Vahlberg from the Biostatistics unit, University of Turku for his advice on statistical analysis. We thank Laboratory of Electron Microscopy at the Institute of Biomedicine, University of Turku. Microscopy and flow cytometry were performed at the Turku Bioscience Cell Imaging and Cytometry core facility, supported by Turku Bioimaging and Euro-Bioimaging consortiums. We thank the personnel for their generous help and expertise. Biocenter Finland is acknowledged for providing the research infrastructures, particularly for cell imaging and cytometry, and electron microscopy.

This manuscript has been released as a Pre-Print at BioRxiv. doi: https://doi.org/10.1101/782276

## 6 Author Contributions

AS and PM designed the study. AS, PP, SHP, VŠ, EK, RV, MV and LC designed and performed the experiments. AS, PP, SHP, LC, MF, YC and PM analyzed the data. PM, YC and MF carried out supervision and provided resources for the project. AS and PM were in charge of writing the manuscript. Additionally, PP, SHP, LC, MF and YC contributed to the text.

## 7 Conflict of Interest

Authors declare no conflict of interest.

## 8 Funding

This work was supported by the Academy of Finland (grant ID: 25700, 296684, 307313, and 327378 to PM; 286712 to VŠ), Sigrid Juselius and Jane and Aatos Erkko foundations (to PM), Turku doctoral programme in molecular medicine (TuDMM) (to SHP and MV). Magnus Ehrnrooth (to AS) and Finnish Cultural (to VŠ) foundations. MF and LC were supported by the Wellcome Trust (212343/Z/18/Z) and EPSRC (EP/S004459/1).

