## Supplementary Figures for "Missing-In-Metastasis / Metastasis Suppressor 1 regulates B cell receptor signaling, B cell metabolic potential and T cell-independent immune responses"

### Supplementary Figure S1

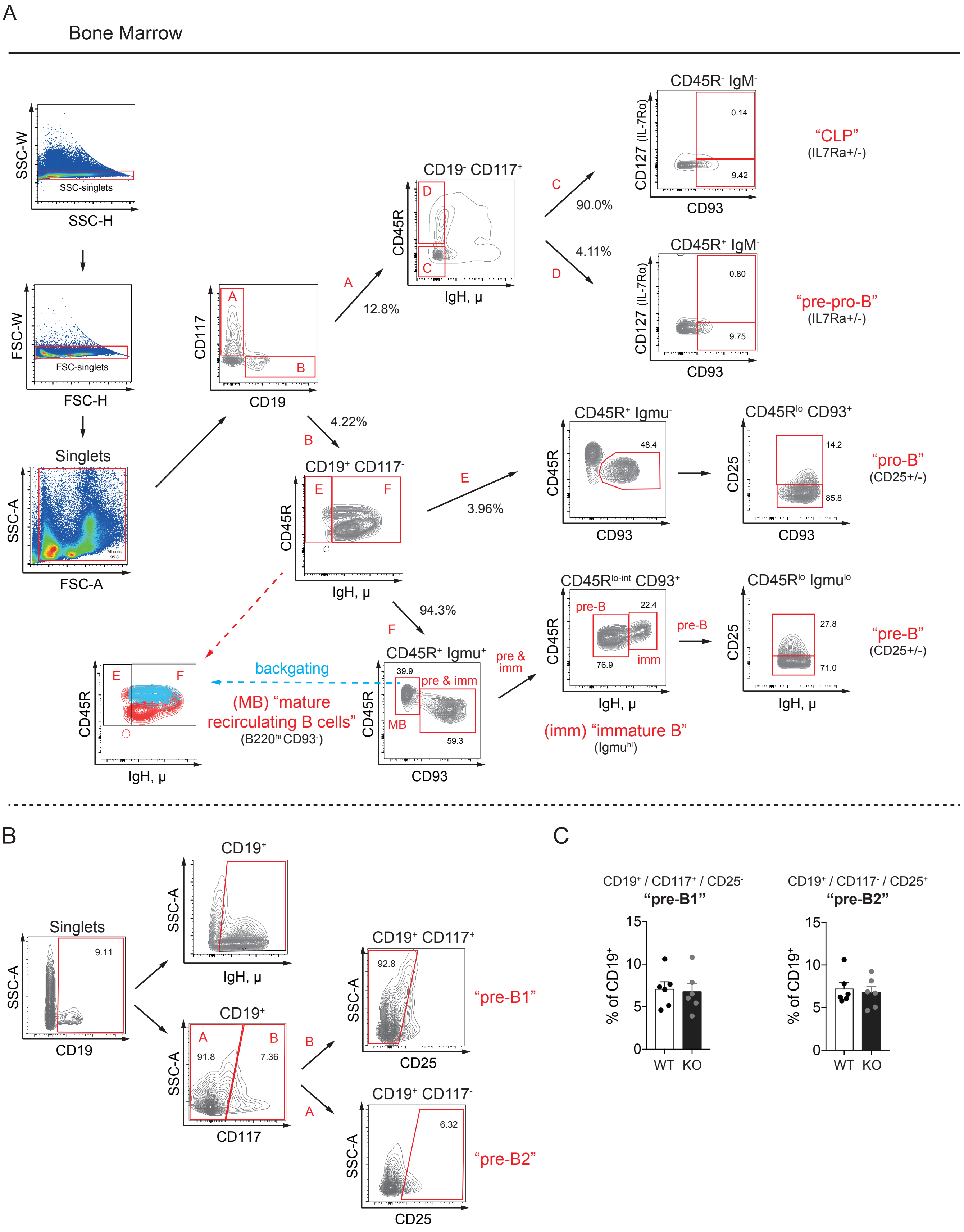


**Supplementary Figure S1. B cell development in the bone marrow of MIM$^{-/-}$ mice.**
**A.** Gating strategy for the analysis of B cell development in the bone marrow (Figure 1D). Bone marrow cell populations are defined as follows. CLP (CD19$^-$ CD117$^+$ / CD45R$^-$ IgM$^-$ / CD93$^+$ CD127$^{+/-}$), pre-pro-B (CD19$^-$ CD117$^+$ / CD45R$^+$+ IgM$^-$ / CD93$^+$ CD127$^{+/-}$), pro-B (CD19$^+$ CD117$^-$ / CD45R$^+$ IgM$^-$ / CD45R$^{lo}$ CD93$^+$ / CD25$^{+/-}$), pre-B (CD19$^+$ CD117$^-$ / CD45R$^+$ IgM+ / CD45R$^{lo\text{-}int}$ CD93$^+$ / CD45R$^{lo}$ IgM$^{lo}$ / CD25$^{+/-}$) immature B (CD19$^+$ CD117$^-$ / CD45R$^+$ IgM$^+$ / CD45R$^{lo\text{-}int}$ CD93$^+$ / CD45R$^{int}$ IgM$^{hi}$), and mature recirculating B cells (CD19$^+$ CD117$^-$ / CD45R$^+$ IgM$^+$ / CD45R$^{hi}$ CD93$^-$). **B.** Gating strategy for the analysis of CD19$^+$ cells in the bone marrow (Figure 1C) and pre-B cells as in Yu et al., 2012. Pre-B1 (CD19$^+$ CD117$^+$ / CD25$^-$), pre-B2 (CD19$^+$ CD117$^-$ / CD25$^+$). **C.** Percentages of pre-B cells in the bone marrow of WT and MIM KO mice as gated in (B). Data of 6 independent experiments. Mean ± SEM.

### Supplementary Figure S2

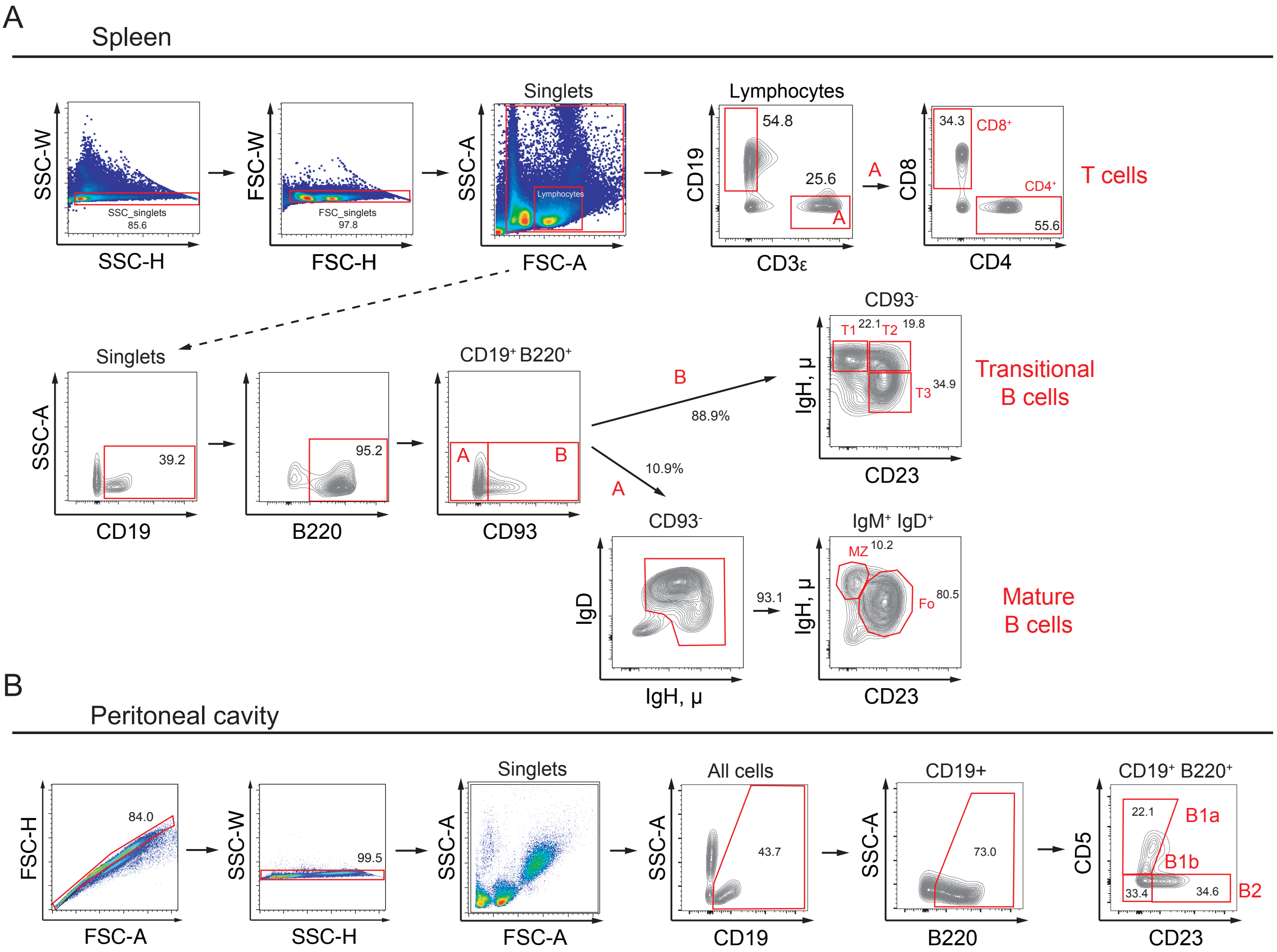


**Supplementary Figure S2. B cell subsets in the spleen and peritoneal cavity of MIM$^{-/-}$ mice.**
**A.** Gating strategy for the analysis of lymphocytes in the spleen (Figure 1E, F). B cells (CD19$^{+}$ CD3ε$^{-}$), CD4$^{+}$ T cells (CD19$^{-}$ CD3ε$^{+}$ / CD4$^{+}$), CD8$^{+}$ T cells (CD19$^{-}$ / CD3ε$^{+}$ / CD8$^{+}$). Transitional B cells – (CD19$^{+}$ / B220$^{+}$ / CD93$^{+}$..) T1 (..CD23$^{-}$ IgM$^{hi}$), T2 (..CD23$^{+}$ IgM$^{hi}$), T3 (..CD23$^{+}$ IgM$^{lo}$). Marginal zone B cells – (MZ, CD19$^{+}$ / B220$^{+}$ / CD93$^{-}$ / IgM$^{+}$ IgD$^{+}$ / CD23$^{-}$ IgM$^{hi}$). Follicular B cells – (Fo, CD19$^{+}$ / B220$^{+}$ / CD93$^{-}$ / IgM$^{+}$ IgD$^{+}$ / CD23$^{+}$ IgM$^{med-hi}$).
**B.** Gating strategy for the analysis of peritoneal cavity B cells (Figure 1G). B1a (CD19$^{+}$ / B220$^{+}$ / CD23$^{-}$ CD5$^{+}$), B1b (CD19$^{+}$ / B220$^{+}$ / CD23$^{-}$ CD5$^{-}$), B2 (CD19$^{+}$ / B220$^{+}$ / CD23$^{+}$ CD5$^{-}$).

Supplementary Figure S3

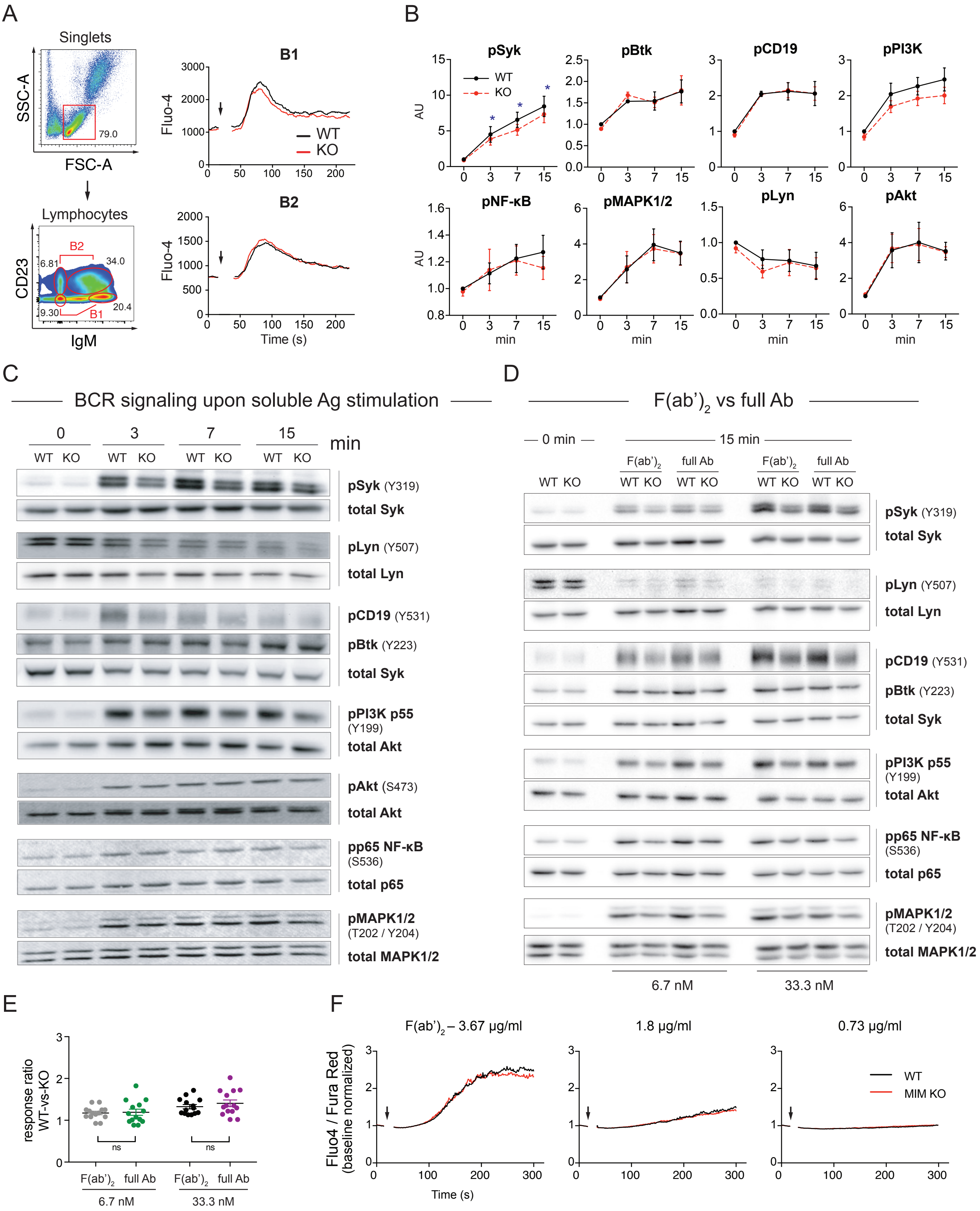

**Supplementary Figure S3. BCR signaling in response to soluble and surface-bound anti-IgM stimulation.**

**A.** Gating strategy and analysis of $Ca^{2+}$ mobilization by peritoneal cavity B cells in response to soluble anti-IgM antibodies (10 μg/ml). Peritoneal cavity B cells were labeled with Fluo-4, pre-stained with anti-CD23-AF593 antibodies to separate B1 and B2 populations, and stimulated with 10 μg/ml of AF647-labeled anti-IgM antibodies. Time of anti-IgM antibody addition is indicated by arrows. Data is presented as Fluo-4 median fluorescence intensity. Representatives of 3 independent experiments are shown. **B-C.** Splenic B cells were stimulated with 5 μg/ml of anti-IgM antibodies in solution for 0, 3, 7 and 15 min and lysed. Lysates were subjected for immunoblotting for phosphorylated forms of different BCR signaling effector proteins. Data is presented as ratios of phosphorylated forms to total protein levels and normalized to the level of WT at 0 min. Data of 4-7 independent experiments. * $p<0.05$. Mean ± SEM. Representative immunoblot images used for quantification are shown. **D.** Comparison of the BCR signaling in response to low and high concentrations of $F(ab')_2$ fragments or full donkey anti-IgM antibodies in WT and $MIM^{-/-}$ B cells. Splenic B cells were stimulated on surfaces coated with 1 (6.7 nM) or 5 μg/ml (33.3 nM) of anti-IgM antibodies or equimolar concentrations of $F(ab')_2$ anti-IgM antibodies for 15 min and lysed. Lysates were subjected for immunoblotting for phosphorylated forms of different BCR signaling effector proteins. Representative immunoblot images of 2 independent experiments. **E.** WT-vs-KO ratios of phosphorylation levels of BCR signaling components were compared between B cells activated for 15 min with surface-adsorbed $F(ab')_2$ and full anti-IgM antibodies at low (6.7 nM) or high (33.3 nM) concentration. Data was pooled from 2 independent experiments and analyzed with repeated measures one-way ANOVA followed by Tukey's multiple comparisons test. **F.** Splenic B cells were labeled as in Fig. 2A and stimulated with $F(ab')_2$ fragment of anti-IgM antibodies in solution at concentrations of 3.67, 1.8 and 0.73 μg/ml (corresponding to molarities of 5, 2.5, and 1 μg/ml of full antibody), and analyzed by flow cytometry. Ratios of Fluo-4 to Fura Red median fluorescence intensity were analyzed and data is presented as mean of 4 independent experiments.

Supplementary Figure S4

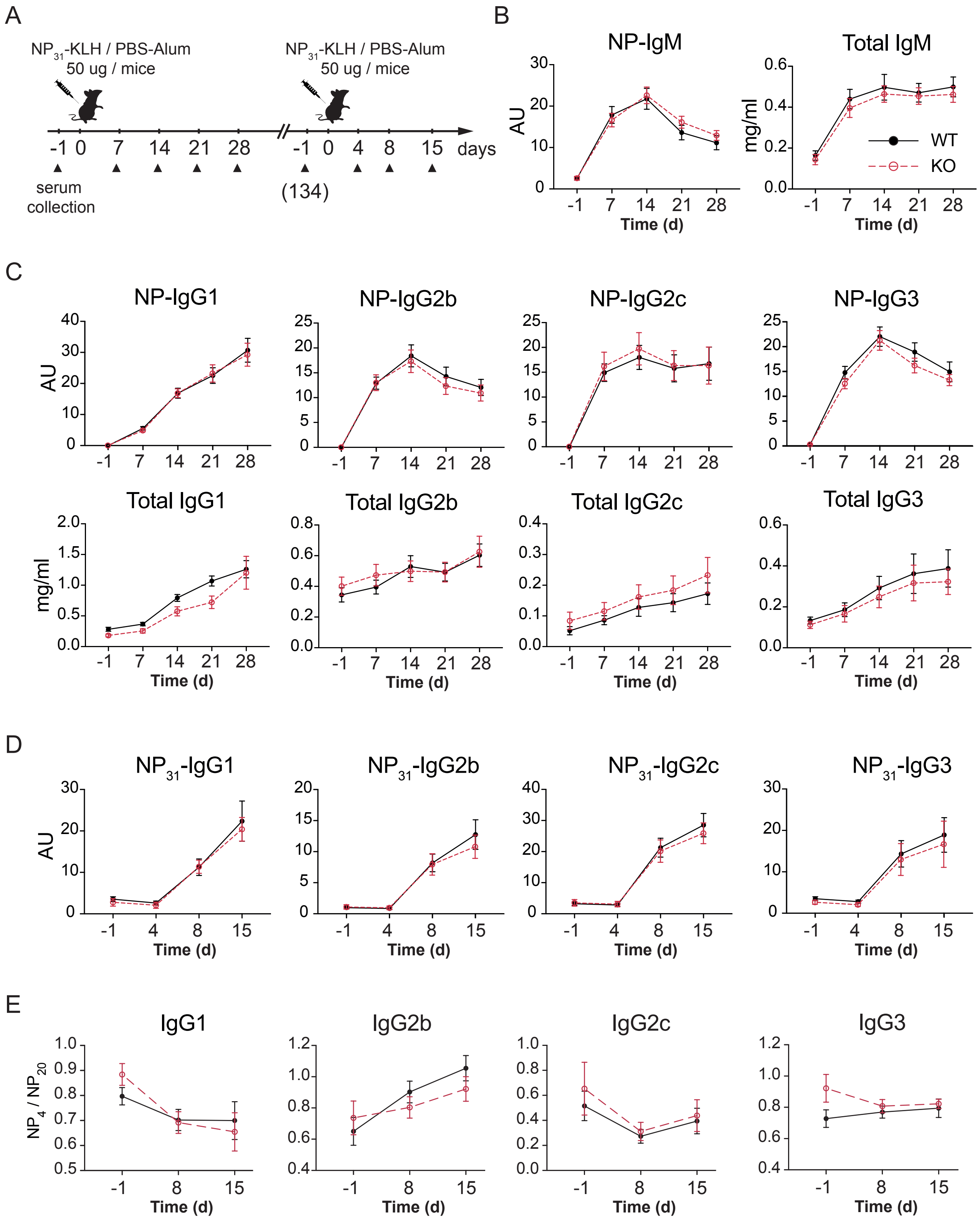


**Supplementary Figure S4. MIM$^{-/-}$ mice mount normal antibody response to T cell-dependent antigen, NP-KLH.**

**A.** A schematic representation of T cell-dependent immunization study.

**B–D.** WT and MIM$^{-/-}$ mice were immunized with NP-KLH as in (**A**). Levels of total and NP-specific antibody levels during the course of T cell-dependent immune response were measured with ELISA. **B.** NP-specific and total IgM antibody levels. **C.** NP-specific and total IgG antibody levels. **D.** NP-specific IgG antibody levels upon secondary immunization with NP-KLH. **E.** Affinity maturation analysis during antibody response upon secondary immunization with NP-KLH. Analysis is based on signal ratio of NP-specific IgG antibodies bound to a haptenated carrier protein in ELISA with low (NP4) and high (NP20) conjugation ratio. 10 mice per group. Mean ± SEM.

Supplementary Figure S5

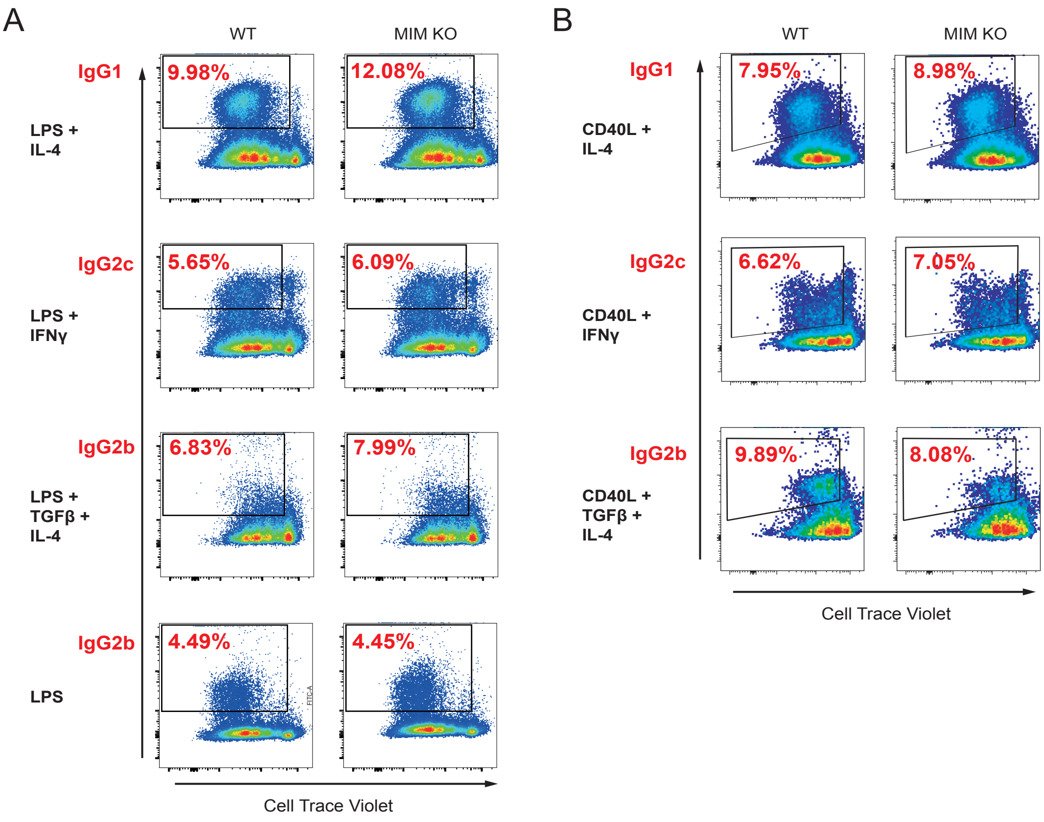


C

| | LPS | | LPS + IL-4 | | LPS + IFNγ | | LPS + TGFβ + IL-4 | |
|---|---|---|---|---|---|---|---|---|
| | WT | KO | WT | KO | WT | KO | WT | KO |
| IgG1 | 0.20 ± 0.13 | 0.19 ± 0.08 | 9.98 ± 2.13 | 12.08 ± 3.07 | 0.14 ± 0.08 | 0.12 ± 0.04 | 1.62 ± 0.57 | 1.85 ± 1.05 |
| IgG2b | 4.49 ± 1.09 | 4.45 ± 0.65 | 8.22 ± 2.16 | 8.99 ± 0.90 | 4.39 ± 0.74 | 4.74 ± 0.80 | 6.83 ± 2.67 | 7.99 ± 2.20 |
| IgG2c | 3.75 ± 1.41 | 3.66 ± 1.42 | 4.03 ± 2.49 | 4.23 ± 2.26 | 5.65 ± 0.81 | 6.09 ± 0.38 | 2.17 ± 1.06 | 2.09 ± 1.19 |
| IgG3 | 4.22 ± 1.99 | 3.86 ± 1.26 | 6.15 ± 4.03 | 5.34 ± 1.68 | 3.72 ± 2.13 | 3.68 ± 1.39 | 4.98 ± 3.13 | 4.79 ± 2.04 |

D

| | CD40L + IL-4 | | CD40L + IFNγ | | CD40L + TGFβ + IL-4 | |
|---|---|---|---|---|---|---|
| | WT | KO | WT | KO | WT | KO |
| IgG1 | 7.95 ± 0.56 | 8.98 ± 1.77 | 0.15 ± 0.05 | 0.16 ± 0.07 | 2.51 ± 0.53 | 3.51 ± 0.48 |
| IgG2b | 8.99 ± 2.08 | 9.36 ± 3.19 | 4.63 ± 1.21 | 4.82 ± 1.56 | 9.89 ± 4.52 | 8.08 ± 4.24 |
| IgG2c | 8.97 ± 2.46 | 9.25 ± 1.65 | 6.62 ± 1.72 | 7.05 ± 2.11 | 8.81 ± 4.47 | 7.95 ± 3.01 |
| IgG3 | 6.54 ± 3.17 | 8.12 ± 4.36 | 2.60 ± 1.05 | 2.93 ± 1.02 | 7.91 ± 4.15 | 7.09 ± 3.93 |

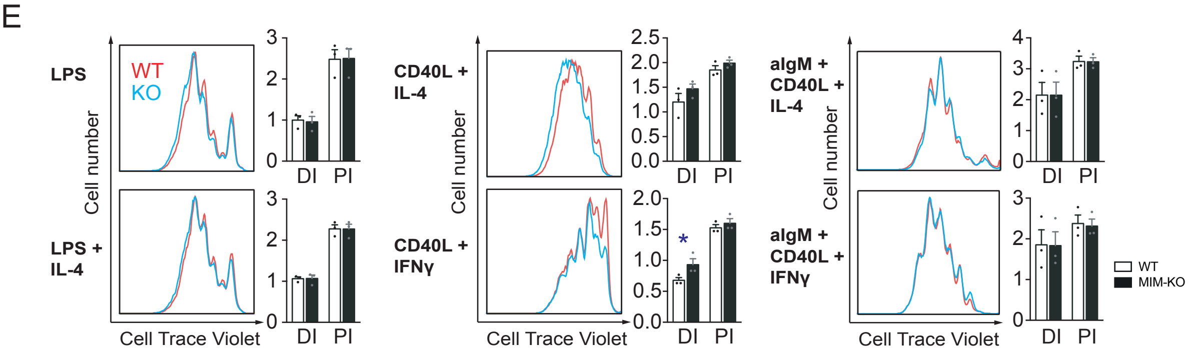


**Supplementary Figure S5. MIM$^{-/-}$ B cells undergo normal class-switching and proliferation in response to LPS, CD40L or IgM-BCR/CD40L stimulation.**

Cell Trace Violet (CTV)-labeled isolated splenic B cells were cultured in LPS- or CD40L-supplemented media with cytokines and analyzed by flow cytometry for surface immunoglobulin expression on day 3. **A.** Analysis of surface IgG expression after 3 day of culture in media supplemented with LPS and cytokines. Representative flow cytometry plots and mean percentages of 3 independent experiments are shown. **B.** Analysis of surface IgG expression after 3 day of culture in media supplemented with CD40L and cytokines. Representative flow cytometry plots and mean percentages of 3 independent experiments are shown. **C.** Results of class-switching experiments from LPS cultures (A). Percentages of IgG$^+$ B cells in each condition are shown. Mean ± SD. **D.** Results of class-switching experiments from CD40L cultures (B). Percentages of IgG$^+$ B cells in each condition are shown. Mean ± SD. **E.** The proliferation of CTV-labeled WT and MIM-KO B cells was analyzed by flow cytometry from cells cultured for 3 days in the presence of LPS alone or with IL-4, CD40L with IL-4 or IFNγ, or surface-adhered anti-IgM in the presence of CD40L and IL-4 or IFNγ. Representative histograms of 3 independent experiments are shown. DI – division index and PI – proliferation index were extracted from the data by FlowJo proliferation platform and are presented as bar graphs. * $p<0.05$. Mean ± SEM.

Supplementary Figure S6

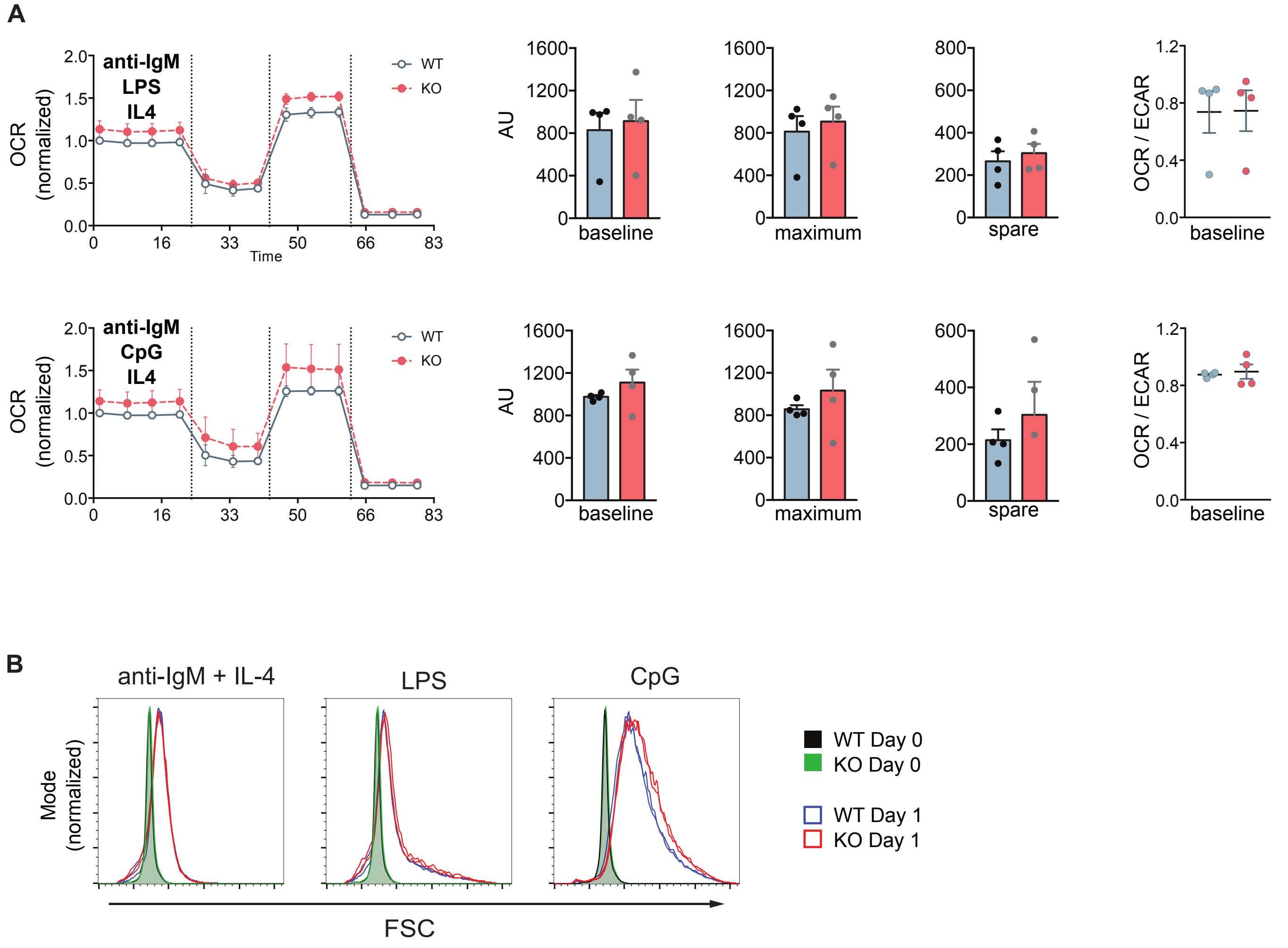


**Supplementary Figure S6. Metabolic activity upon stimulation with Toll-like receptor ligands LPS or CpG in combination with BCR.**
**A.** Oxygen consumption rate (OCR) profiles of WT and MIM-KO splenic B cells stimulated with: anti-IgM + LPS + IL-4 or anti-IgM + CpG + IL-4 for 24 h were measured in a Seahorse XF Cell Mito Stress Test assay. The different steps of the assay are depicted in the graph on the left showing mean ± SEM. Comparisons of baseline mitochondrial respiration as well as maximum and spare respiratory capacities, extracted from the assay, are shown in the graphs in the middle, and quantification of the ratio of OCR to ECAR (extracellular acidification rate) at the baseline is shown on the right. Data is from 4 independent experiments. Mean ± SEM. **B.** Comparison of WT and MIM-KO splenic B cells size prior (day 0) and after 24h of stimulation (day 1) with: anti-IgM + IL-4, LPS or CpG, measured by flow cytometry as forward scatter intensity. Representative data of 6 independent experiments.
